# A Perspective on Twenty Years of Human-Leopard Conflict in Pauri Garhwal, Uttarakhand, India with Sustainable Solutions for its Mitigation

**DOI:** 10.1101/2025.08.15.669212

**Authors:** S. M. Chowfin

**Affiliations:** C/o The Gadoli and Manda Khal Wildlife Conservation Trust, P.O. Box 27, Pauri, District Pauri Garhwal, Uttarakhand, India

**Keywords:** Anthropogenic pressures, central Himalayas, ecological restoration, human – leopard conflict, Pauri Garhwal, *Van Panchayats*

## Abstract

Human - leopard conflict poses a significant conservation challenge in the Pauri Garhwal district of Uttarakhand, India, with 290 reported human – leopard conflict incidents between 2000 and 2020, averaging 13.81 incidents annually. The most affected age group were adults followed by children with a noticeable increase in incidents in winter. Key factors driving this conflict include habitat degradation, loss of prey, and human encroachment on forest areas. Interestingly, the Gadoli and Manda Khal Fee Simple Estates, an area undergoing ecological restoration, by forest protection through foot patrols, fire watches and passive forest restoration (natural regeneration) within this human – leopard conflict hotspot, reported no incidents during the same timeframe, highlighting the positive impact of habitat-focused interventions for mitigating human – leopard conflict. In contrast, reactive measures such as lethal control lack precision, targeted protocols and long-term efficacy. This study emphasizes ecological restoration as a sustainable strategy for mitigating conflict and calls for improved monitoring of leopard populations, prey species, and habitat conditions. It also offers recommendations for effective conflict management in the region, stressing the advantages of restoration over reactive approaches.

## 1. Introduction

Human-carnivore conflict is a pervasive challenge in biodiversity conservation, with profound ecological and socio-economic implications. Across the globe, interactions between large carnivores and human populations often result in negative outcomes, including livestock depredation, human injuries, and fatalities, which fuel local animosity toward wildlife [Treves & Karanth, 2003; Inskip & Zimmermann, 2009]. These conflicts, if unmanaged, jeopardize both human safety and the conservation of threatened species [Woodroffe et al., 2005]. In the Indian subcontinent, the Indian leopard (*Panthera pardus fusca*) represents a notable case of human-carnivore conflict. As a highly adaptable carnivore, leopards occupy diverse habitats, from dense forests to human-dominated landscapes, surviving on a broad dietary spectrum ranging from wild ungulates to livestock and, in some cases, human prey [Jacobson et al., 2016; Naha et al., 2020]. However, this adaptability also brings leopards into frequent conflict with humans, particularly in regions where their natural habitats are degraded or fragmented [Athreya et al., 2011; Stein et al., 2020; Stein et al. 2024].

### 1.1 Background

Human-carnivore conflicts arise from threats to human life, economic security, or on outdoor pursuits and recreational activities caused by the action of carnivores and are a significant issue where humans and carnivores coexist. These conflicts can involve competition over resources and space, often resulting in predation, property damage, or injuries to livestock, pets, or humans [Treves & Karanth, 2003; Woodroffe et al., 2005; Klees van Bommel et al., 2020]. In the Indian Himalayan Region (IHR), particularly in the Pauri Garhwal district of Uttarakhand, leopard-related conflicts have become a growing conservation issue. While livestock predation is a notable concern, leopard attacks on humans have escalated, fueling negative attitudes and increasing calls for extermination [Naha et al., 2018; *pers. Obs*.]. Several factors contribute to the intensification of human-carnivore conflict, including habitat loss, fragmentation, reduced prey availability, and increasing human populations near wilderness areas [Woodroffe, R., 2000; Treves & Karanth, 2003; Jacobson et al., 2016; Parchizadeh & Belant, 2021]. Despite the ending of eradication programs in most areas, certain regions continue to engage in lethal control, such as hiring private hunters or using trained agents to eliminate problematic carnivores [Treves & Karanth, 2003]. In Uttarakhand’s mountainous regions, human – leopard conflict is the primary type of human-wildlife conflict, and the management of human-leopard conflict in Uttarakhand is largely centered around lethal control, where leopards responsible for human harm or fatalities are targeted for elimination. However, the approach is flawed, as there is no established protocol for identifying problematic leopards within the leopard population. The decision to remove a leopard is left to the discretion of private hunters or department staff, making targeted control challenging [Athreya et al. 2011]. In addition to lethal control, leopards are sometimes trapped or tranquilized (Fig. 1) by the Uttarakhand Forest Department, and are often mistreated and traumatized (Fig.2.) (Video File 1: https://doi.org/10.6084/m9.figshare.23620626.v1) but, as with lethal measures, there is no prior identification process to ensure that the correct leopard individual is identified. The current management strategies, particularly lethal control and trapping, do not adequately address the socio-ecological dynamics of human-leopard conflicts, necessitating the development of a more targeted, humane, and scientifically informed approach to mitigate these conflicts.

**Figure 1.**
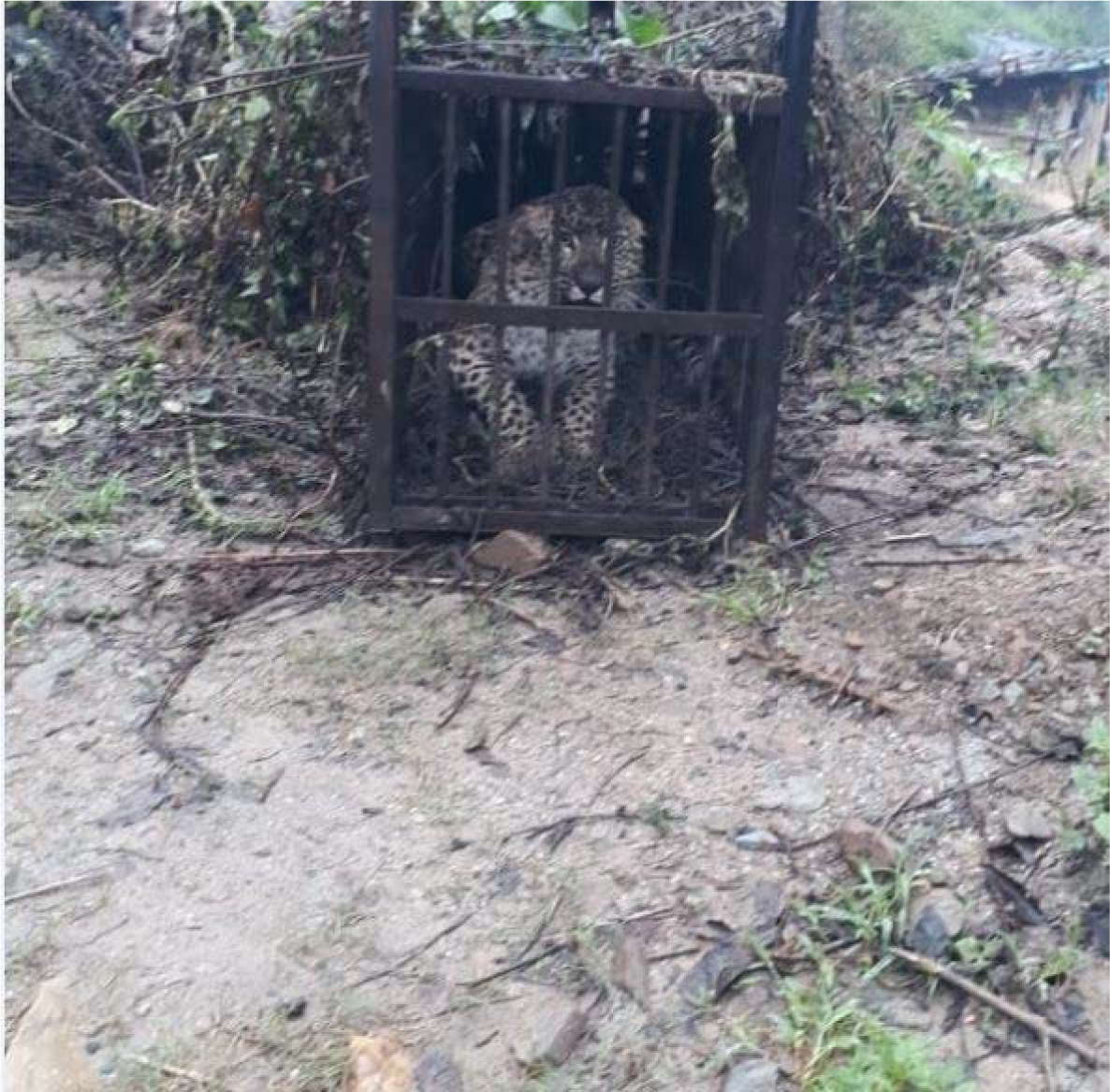
A leopard (Panthera pardus fusca) trapped by the Uttarakhand Forest Department, Pauri Garhwal, Uttarakhand, India

**Figure 2.**
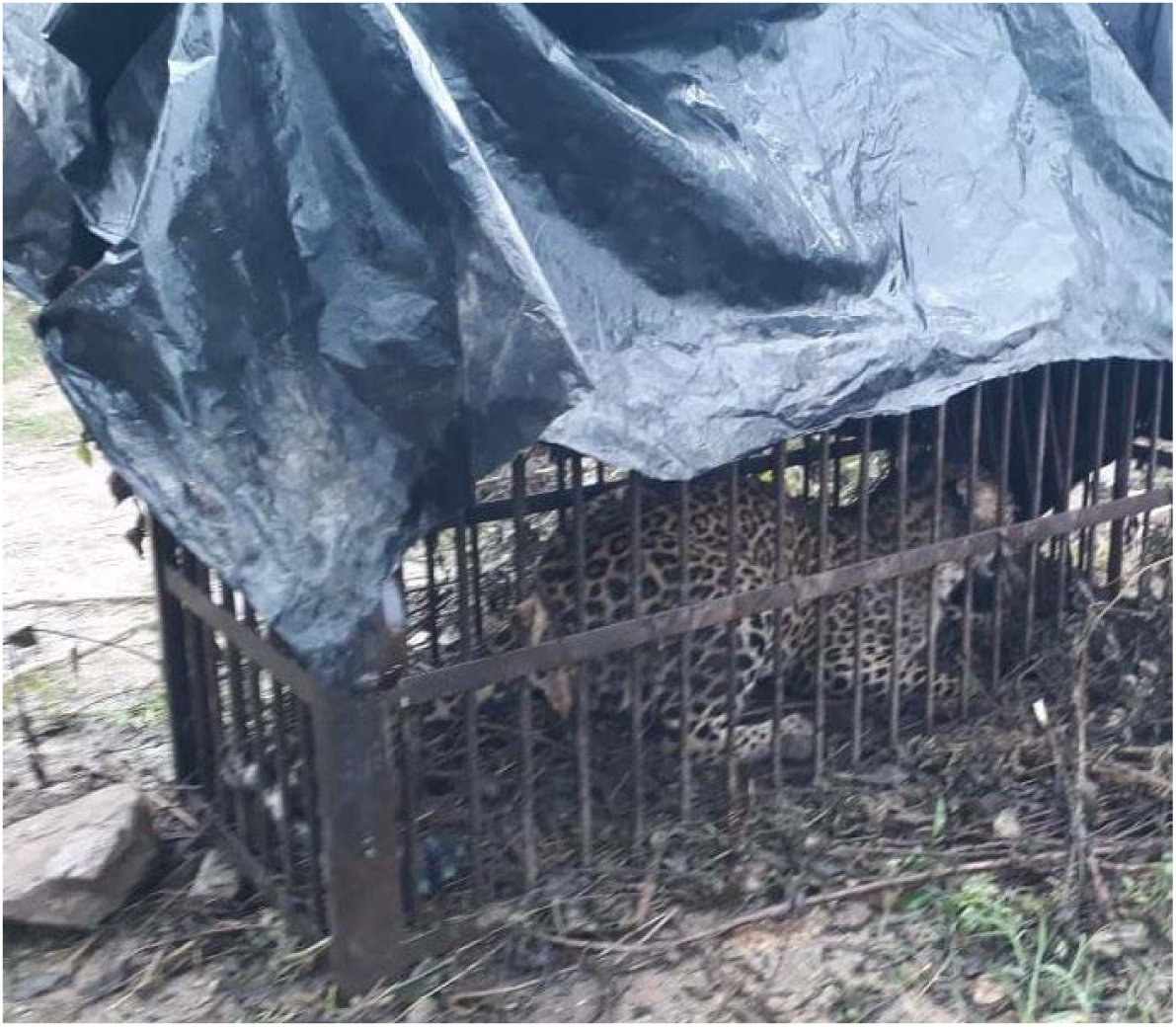
A leopard (Panthera pardus fusca) trapped by the Uttarakhand Forest Department being traumatized during the process (Pauri Garhwal, Uttarakhand, India)

### 1.2 Study Context

The Pauri Garhwal district in Uttarakhand, India, exemplifies the challenges of human-leopard coexistence. From 2000 to 2020, the district witnessed 290 reported human – leopard conflict incidents, averaging 13.81 incidents annually. These incidents are symptomatic of broader ecological issues, including habitat fragmentation, prey depletion, and human encroachment into forested areas [Mina et al., 2023]. The annual persistence of such conflict highlights the need for sustainable mitigation strategies that address the underlying drivers rather than relying solely on reactive measures such as lethal control [Karanth & Chellam, 2009]. Despite the prevalence of conflicts, few studies have examined the effectiveness of mitigation measures in regions like Pauri Garhwal. Reactive approaches, such as trapping or culling, often lack targeted protocols and fail to address the ecological and social dimensions of the problem [Treves et al., 2006]. In contrast, ecological restoration initiatives, such as those implemented in the Gadoli and Manda Khal Fee Simple Estates [Chowfin & Leslie, 2021], show promise by reducing human-wildlife interactions; and creating conditions for leopard prey by habitat recovery. These estates, located in a human – leopard conflict hotspot, reported zero leopard attacks during the two decades studied, underscoring the potential of long-term, proactive habitat conservation efforts.

### 1.3 Objectives

Against this background, this study was conducted to:

- Analyze patterns and trends in human-leopard conflict in Pauri Garhwal over two decades.
- Identify the primary drivers of conflict, including ecological and anthropogenic factors.
- Evaluate the effectiveness of existing mitigation strategies, contrasting reactive measures with proactive habitat restoration.
- Provide actionable recommendations for sustainable conflict mitigation in the region.

By integrating ecological, social, and management perspectives, this study aims to contribute to the growing body of knowledge on human-carnivore coexistence and inform evidence-based conservation policies.

## 2. Drivers of conflict

The major drivers underlying human – leopard conflict in the district as per this study were infrastructure development in forest areas, human - induced forest fires, resource extraction which included illicit felling and lopping and illegal grazing of cattle leading to deforestation and degradation of leopard prey habitat, the alienation of *Van Panchayats* and dumping of garbage in the open in villages and reserve forests.

### 2.1. Development in forest areas

Development in forest areas of the Pauri Garhwal district mainly involved 5 types of infrastructure projects viz. power lines, roads, buildings, railways and quarrying. Community-managed forests were exploited more than reserve forests, with 87.9% (n=713.5879 ha/812.0915 ha) of forest area used for infrastructure being community-managed forests and 12.1% (n=98.5126 ha/812.0915 ha) being forest department-managed reserve forest. Power lines constituted 51.8% (n = 420.9276 ha/812.0915 ha) of the forest area exploited, roads made up 41.8% (n = 339.4318 ha/812.0915 ha), buildings made up 3.6%, railways made up 2.38% and quarrying made up 0.37% of the forest area exploited in the Pauri Garhwal district (Table 1) which are conservative lower bound estimates.

**Table 1.** Infrastructure projects (ha) in forests in Pauri Garhwal (2000 – 2020) (sourced from the Uttarakhand Forest Department – Pauri under the Right to Information Act, 2005)

| **Infrastructure Project** | **Area used in Reserve Forest** | **Area used in Community Forest** | **Total** |
| --- | --- | --- | --- |
| Power lines | 17.3071 | 403.6196 | **420.9267** |
| Roads | 75.7505 | 263.6813 | **339.4318** |
| Buildings | 5.455 | 23.899 | **29.354** |
| Quarry | 0 | 3.03 | **3.03** |
| Railway | 0 | 19.349 | **19.349** |
| **Total** | **98.5126** | **713.5789** | **812.0915** |

### 2.2. Forest fires

The period from 2002-2019 saw an estimated 17831 incidents of forest fires in Pauri Garhwal with an average of 990.6□±□884.9 (mean ± sd) forest fires per year [Mina et al., 2023]. In the Pauri sub-division, between 2000 and 2019, an estimated 1423 forest fires occurred with an average affected (i.e. burn) area of 3.36 ha. 74.8% (n=1065/1423) occurred in community-managed forests and 25.2% (n=358/1423) occurred in forest department-managed reserve forests (Table 2). These are conservative lower bound estimates as many forest fires go undetected and unreported.

**Table 2.** Forest fires in the Pauri Sub-division (2000 – 2019) (sourced from the Uttarakhand Forest Department – Pauri under the Right to Information Act, 2005)

| **Year** | **Incidents in Reserve**<br>**Forests** | **Incidents in Community**<br>**Forests** | **Total number of**<br>**incidents** | **Average size of**<br>**affected area (ha)** |
| --- | --- | --- | --- | --- |
| 2000 | 0 | 29 | 29 | 3.91 |
| 2001 | 2 | 44 | 46 | 8.5 |
| 2002 | 11 | 25 | 36 | 3.02 |
| 2003 | 7 | 0 | 7 | 7.36 |
| 2004 | 21 | 11 | 32 | 4.36 |
| 2005 | 15 | 7 | 22 | 6.09 |
| 2006 | 0 | 0 | 0 | 0 |
| 2007 | 3 | 12 | 15 | 7.82 |
| 2008 | 18 | 22 | 40 | 4.15 |
| 2009 | 21 | 64 | 85 | 5.06 |
| 2010 | 5 | 28 | 33 | 3.86 |
| 2011 | 3 | 8 | 11 | 1.91 |
| 2012 | 47 | 57 | 104 | 2.41 |
| 2013 | 4 | 34 | 38 | 1.36 |
| 2014 | 17 | 42 | 59 | 1.26 |
| 2015 | 12 | 28 | 40 | 1.59 |
| 2016 | 34 | 166 | 200 | 2.29 |
| 2017 | 4 | 78 | 82 | 1.44 |
| 2018 | 121 | 323 | 444 | 3.97 |
| 2019 | 13 | 87 | 100 | 2.25 |
| **Total** | **358** | **1065** | **1423** | **3.36** |

### 2.3. Resource Extraction

Illicit felling of natural regeneration of trees is a severe threat to natural forests in the landscape. Prized trees include *Quercus* spp., kafal (*Myrica esculenta*) and other broadleaf evergreen tree species (Fig.3.). In addition to illicit felling; lopping and illegal grazing of cattle in forests drive forest degradation and deforestation in the landscape. Though instances were not quantified in this study several observations were made of these activities in forests.

**Figure 3.**
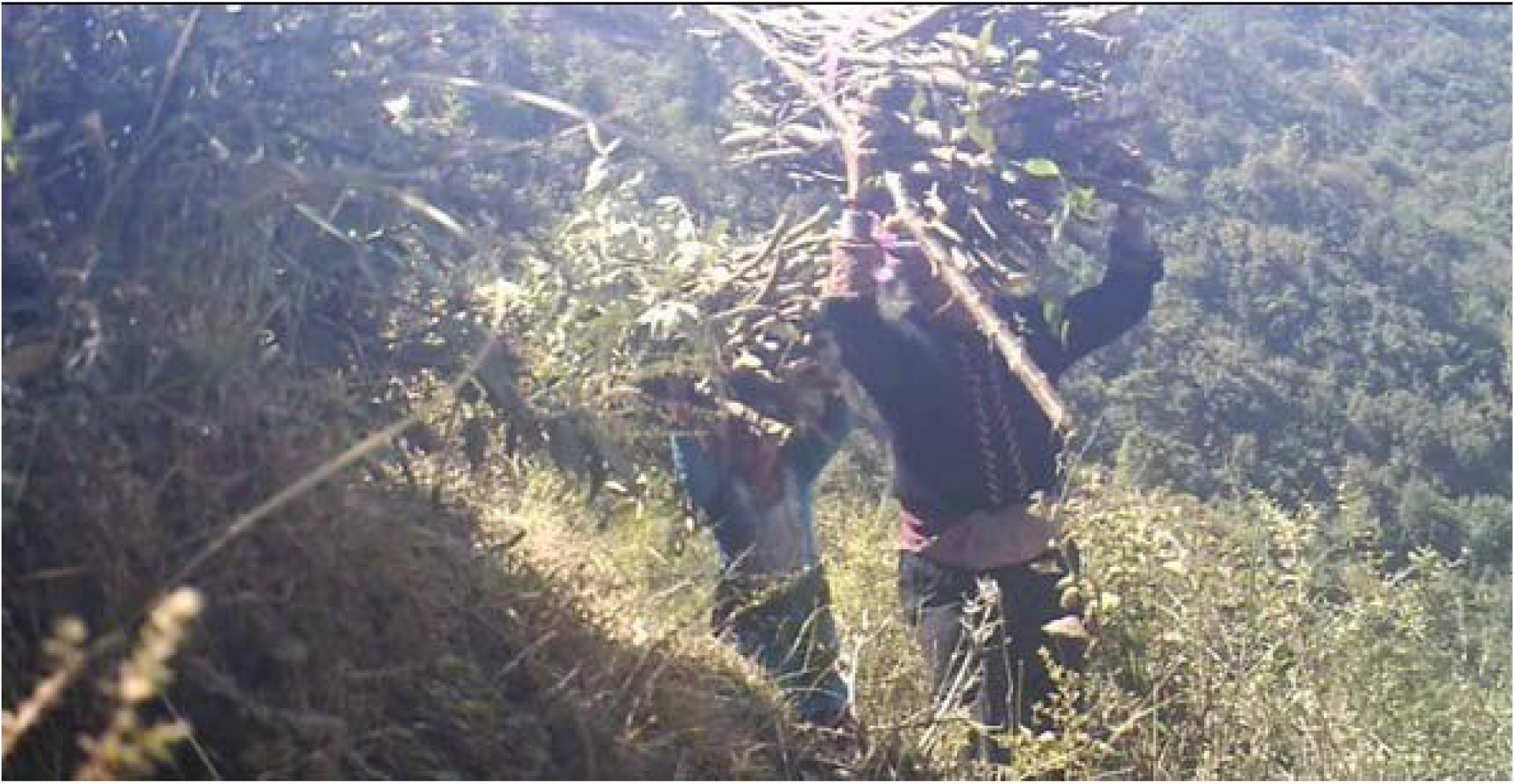
Illicit felling of broadleaf species (Pauri Garhwal, Uttarakhand, India)

### 2.4. Alienation of *Van Panchayats*

During the Colonial Period, British officials acknowledged the role of local communities in forest management based on the Kumaon Forest Grievances Committee’s recommendations. The Forest Panchayat Act of 1931, enacted under Section 28(2) of the Indian Forest Act, 1927, established village forest councils (*Van Panchayats*) with authority to regulate forest resource access, monitor usage, enforce penalties, and manage income for forest welfare. Currently, Uttarakhand has 12,167 Van Panchayats managing 7,326.889 km² of forest area [Naaz & Sahu, 2018], including 2,165 in Pauri Garhwal district covering 386.5141 km² [UKFD, 2023]. Post-independence amendments to the Van Panchayat Act in 1976, 2001, 2005, 2012 [Naaz & Sahu, 2018] and 2024 (*pers. Obs.*) progressively centralized decision-making, reducing community control and increasing bureaucratic oversight. These changes have led to the alienation of *Van Panchayats* from managing community forests [Naaz & Sahu, 2018].

### 2.5. Garbage dumps

Dumping of garbage in the open was observed in several villages sharing a common interface with forests including reserve forests but instances were not quantified in this study.

## 3. Broader Perspectives

Globally, human-carnivore conflicts share common drivers, including habitat loss, prey depletion, and human encroachment. Comparisons with other systems, such as lion-human conflicts in Africa [Loveridge et al., 2007] or cougar-human conflicts in the Americas [Lindzey et al., 1992], reveal valuable insights. For example, community-based conservation initiatives in Kenya [Western et al., 2015] have demonstrated success in reducing conflict while fostering local stewardship. Incorporating lessons from these contexts can enhance the applicability and effectiveness of mitigation strategies in Pauri Garhwal.

## 4. Methods

### 4.1 Study Area

The study area is the Pauri Garhwal district (Fig.4) located in the western Himalayan state of Uttarakhand, India with an area of 5,329 square kilometers. It is situated between 29° 45’ to 30°15’ North and 78° 24’ to 79° 23’ East.

**Figure 4.**
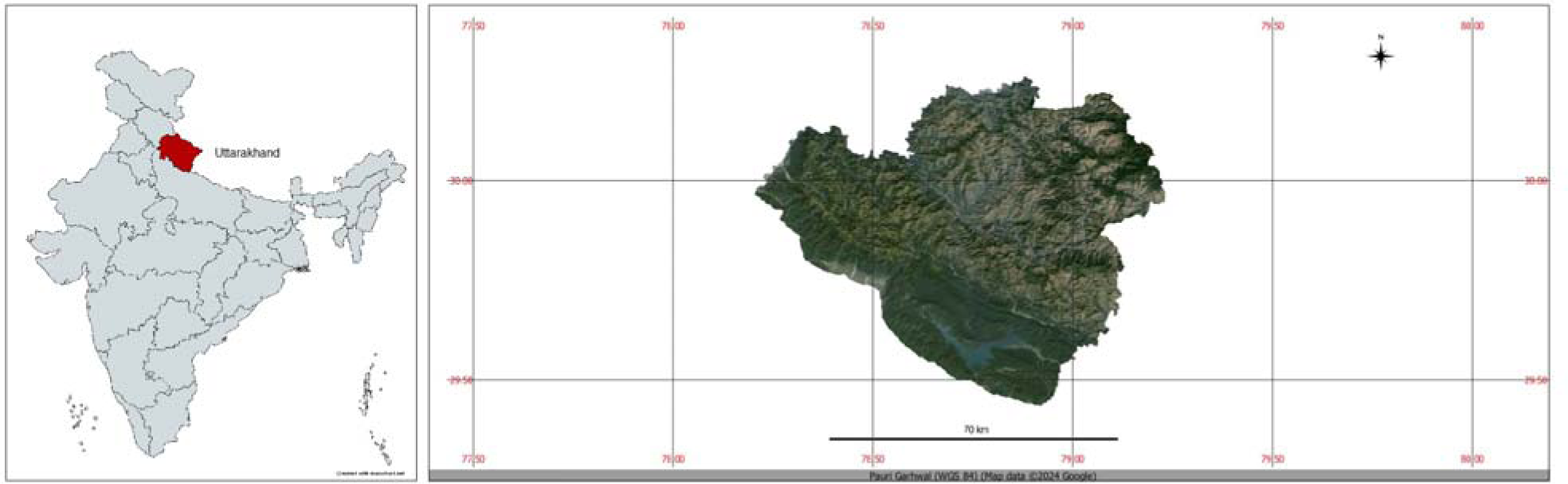
Map of the Pauri Garhwal District in Uttarakhand, India

The total population in the district is estimated to be 687,271 persons with 360,442 females and 326,829 males at a sex ratio (per 1000 persons) of 1103 females per 1000 males and a population density of 129 persons / km^2^. The climate in the district varies from sub-temperate to temperate depending on elevation. The average annual rainfall in the district is 218 cm concentrated during the monsoon period though rain and snowfall also occur in winter. The average temperature ranges between 24 C to 30 C with the maximum temperature being 45 C and the minimum temperature being 1.8 C. The Pauri Garhwal district is divided into six sub-divisions for administrative purposes viz. Pauri, Srinagar, Lansdowne, Thalisain, Dhumakot and Kotdwar. [DEIAA, 2018]. For management of forests, the forests in the district are largely administered by the Garhwal Forest Division of the Uttarakhand Forest Department. Forest cover in the district is estimated to be 3396.71 square kilometers representing 63.74% (n=3396.71/5329) of its geographical area [FSI, 2021]. Forests are managed as protected areas, forest department-managed reserved forests also known as territorial forests, community forests managed by *Van Panchayats*, Civil Soyyam forests or as Private Forests [Singh, 2013; Somanathan, 1991; Chowfin, 2016; Chowfin & Leslie, 2021].

### 4.2 Conflict - Mitigation site

The conflict - mitigation site defined as the site where human – leopard conflict mitigation measures are being implemented and to which human – leopard conflict is being compared to, encompasses the privately owned Gadoli and Manda Khal Fee Simple Estates, covering approximately 4.5 km² of private forest and some agricultural land, with no third-party rights. Located in the central himalayan tracts of the Pauri Garhwal district, Uttarakhand, India, the estates lie at Latitude 30° 07’ 21” N and Longitude 78° 48’ 12” E [Chowfin, 2016; Chowfin & Leslie, 2021] within a human-leopard conflict hotspot identified by Agarwal [2011]. Leopards (*Panthera pardus fusca*) (Fig.5) are resident on these estates.

**Figure 5.**
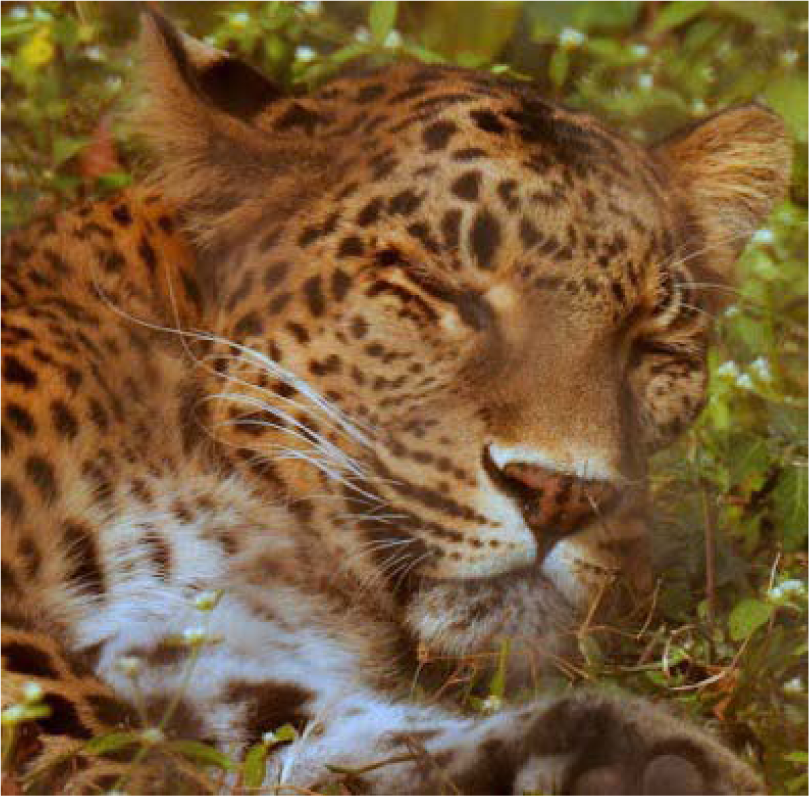
A Leopard (Panthera pardus fusca) in the Gadoli and Manda Khal Fee Simple Estates, Pauri Garhwal, Uttarakhand, India

and are monitored via camera traps (Fig. 6a & 6b) (Video Files 2a & 2b: https://doi.org/10.6084/m9.figshare.23620626.v1) and sign surveys.

**Figure 6a & 6b.**
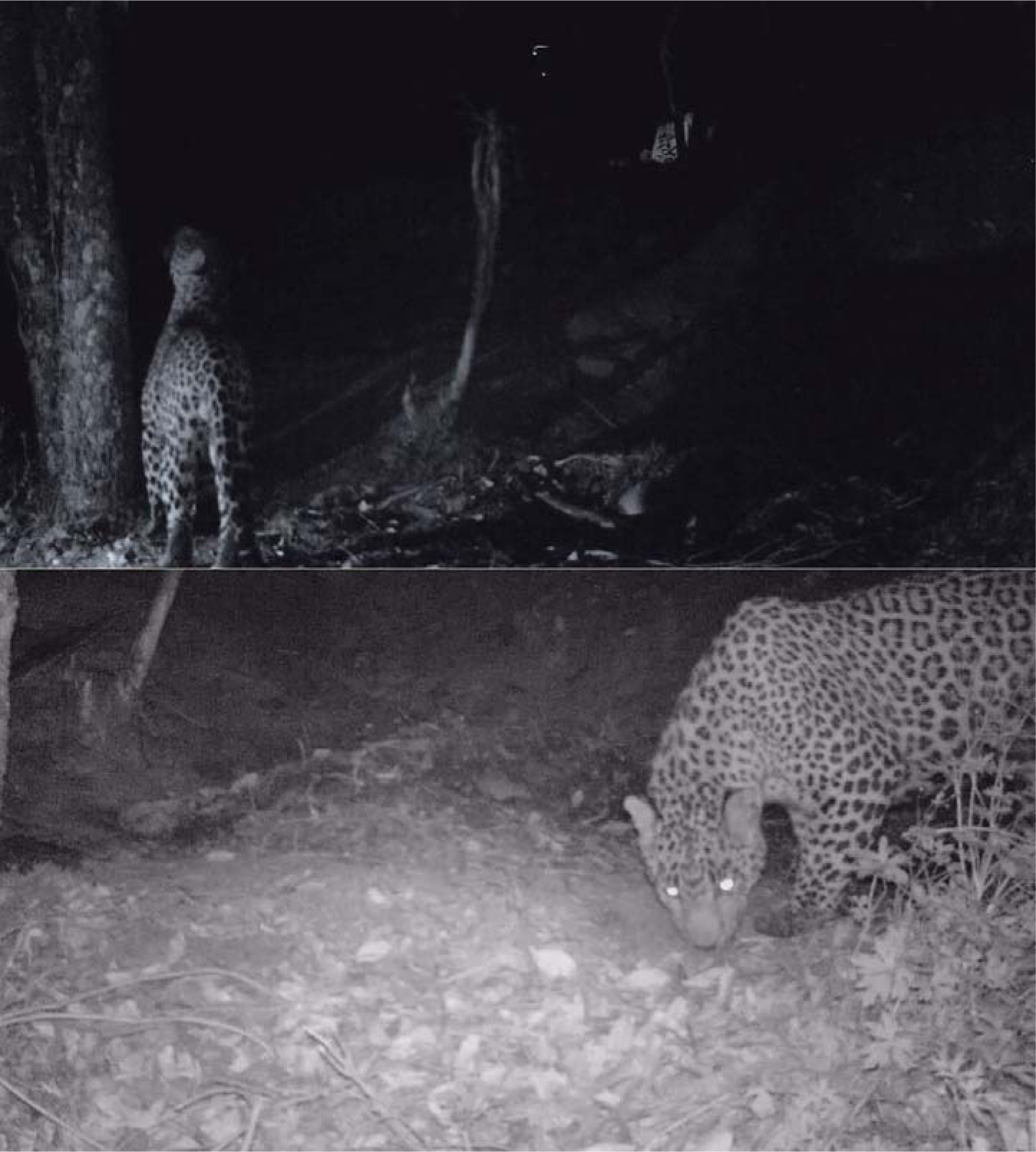
Monitoring of leopards with camera – traps in the Gadoli and Manda Khal Fee Simple Estates, Pauri Garhwal, Uttarakhand, India.

Prey species are also monitored through camera traps (Video Files 3a & 3b: https://doi.org/10.6084/m9.figshare.23620626.v1), visual encounter surveys, and sign surveys. Several conservation actions have been undertaken to protect and restore the private forests of the Gadoli and Manda Khal Fee Simple Estates, including legal measures to address illegal non-forest activities [Chowfin & Leslie, 2021]. Human-leopard conflict mitigation efforts focus on ecological restoration, aiming to (i) reduce human disturbances to allow leopard and prey habitats to recover, and (ii) prevent degradation of undisturbed forests (Fig.7).

**Figure 7.**
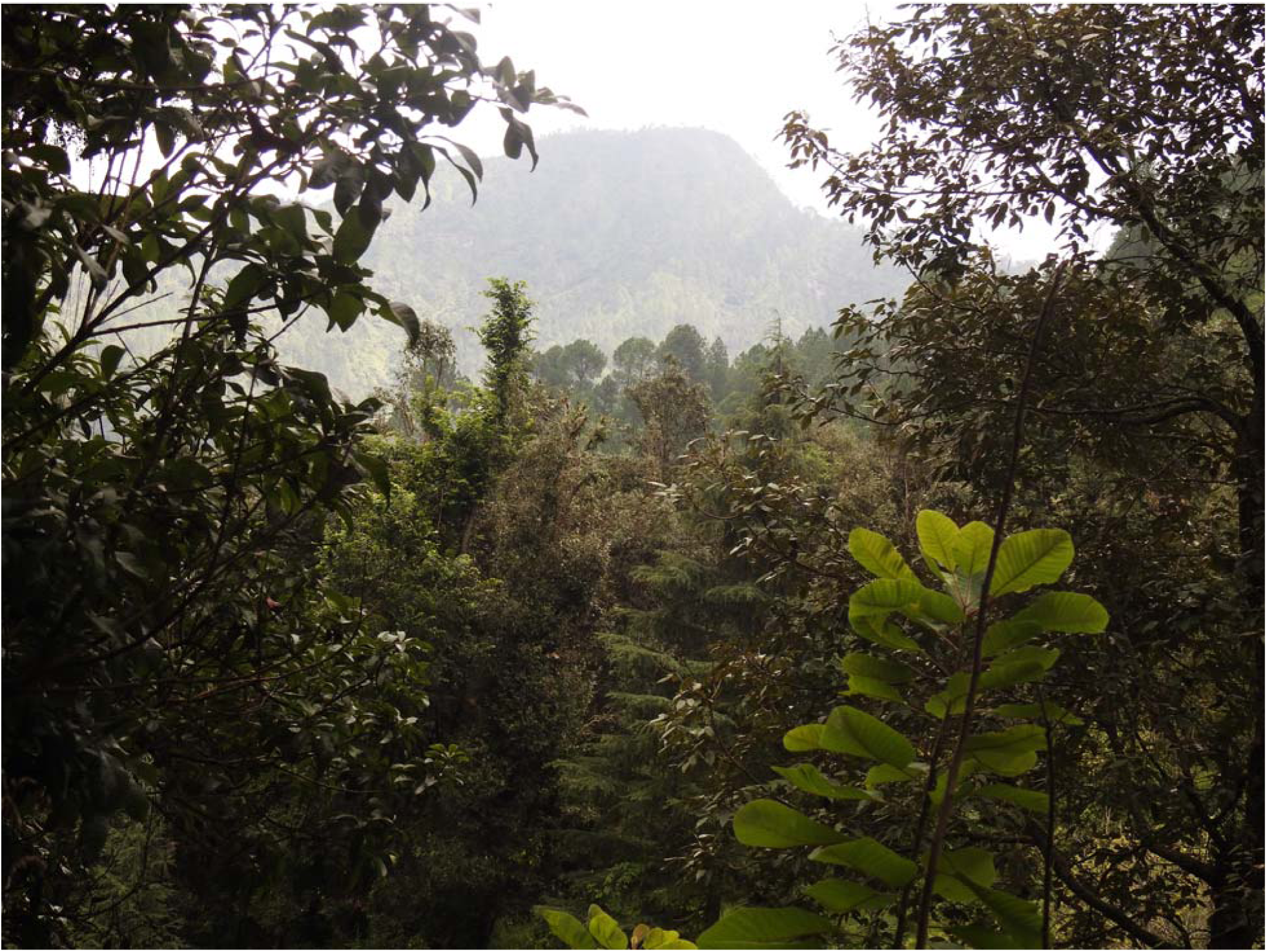
Undisturbed forest tracts in the Gadoli and Manda Khal Fee Simple Estates, Pauri Garhwal, Uttarakhand, India

This approach hypothesizes that with lesser human disturbances to these forests, the quality and extent of forest habitat would improve and increase, creating conditions conducive for leopard prey. This in turn, would curtail leopard attacks on people on these estates, due to the resulting presence of leopard prey, reducing the need for leopards to enter into conflict with people resident on these estates. Human population density in the estates is relatively low compared to the Pauri Garhwal district (129 persons/km²) and Pauri subdivision (177 persons/km²) [DEIAA, 2018] but similar to surrounding villages.

Since 2016, forest patrols conducted at least thrice weekly have curtailed human entry, apprehending trespassers involved in activities like illicit lopping (Fig. 8), cutting forest regeneration (Fig.9), setting forests on fire in nearby areas (Fig.10) (Video Files 4a & 4b: https://doi.org/10.6084/m9.figshare.23620626.v1), and illegal grazing, all of which degrade forest habitat.

**Figure 8.**
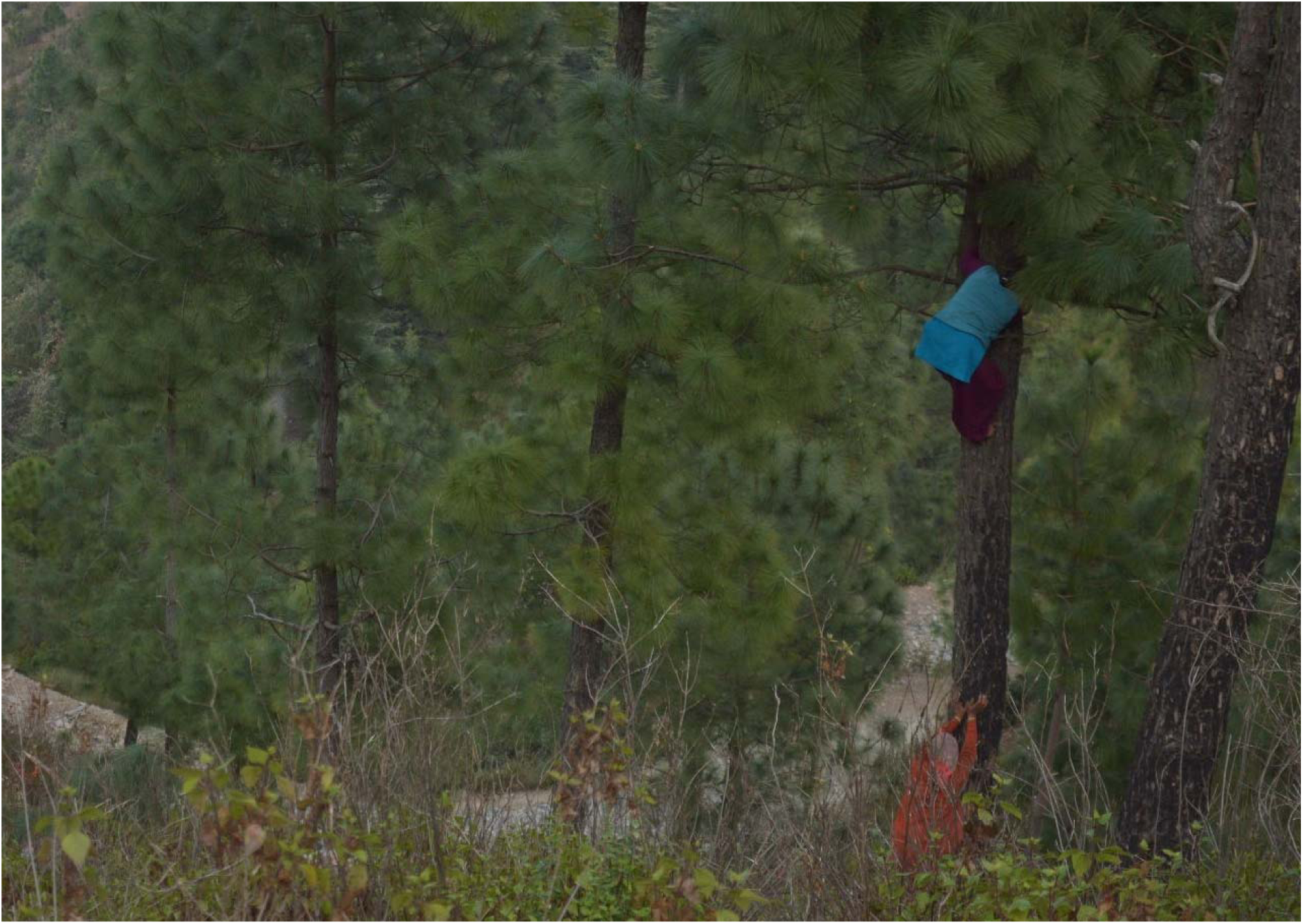
Trespassers being apprehended by the forest patrol while lopping the forest canopy in the private forests of Gadoli and Manda Khal, Pauri Garhwal, Uttarakhand, India

**Figure 9.**
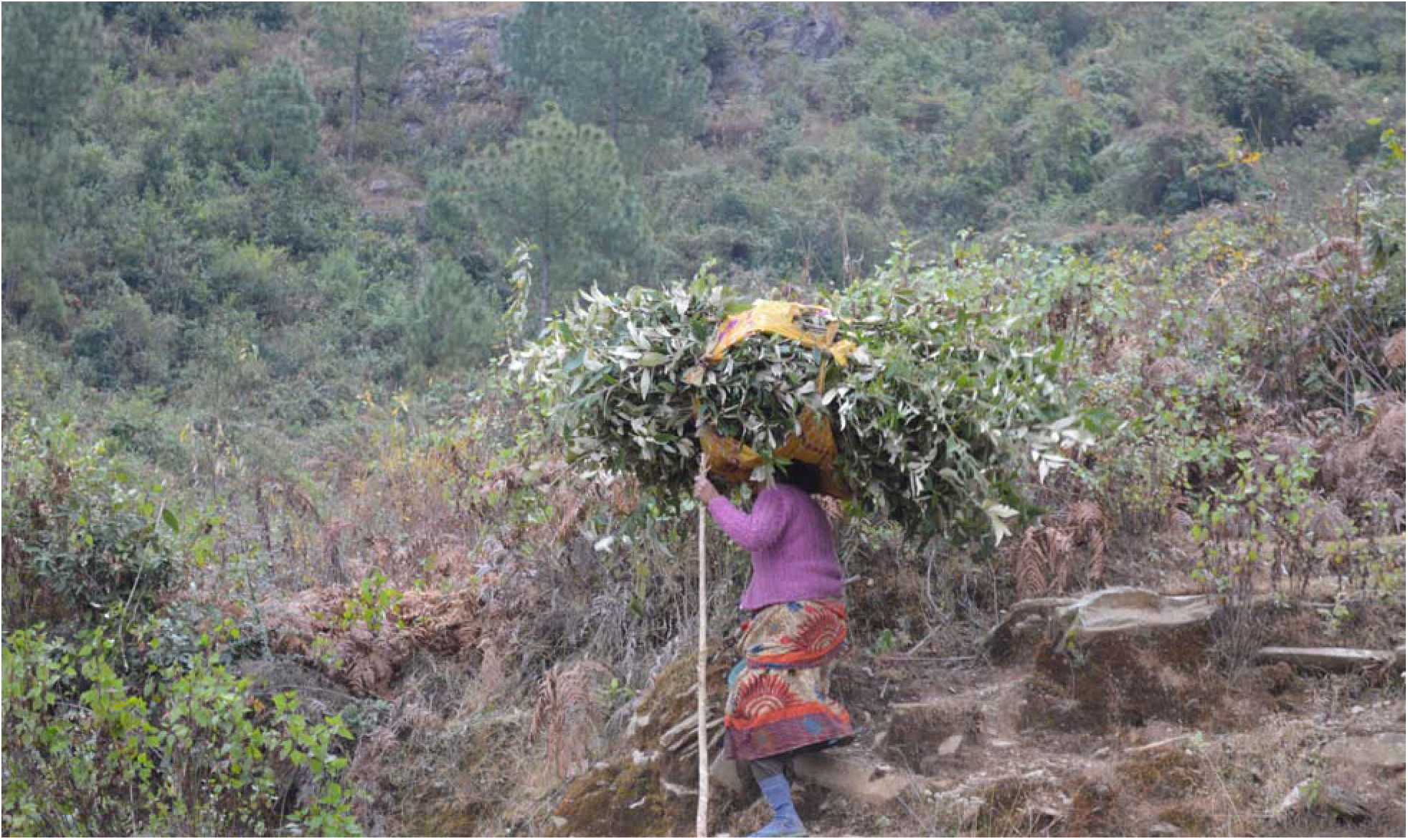
A trespasser being apprehended by the forest patrol while cutting forest regeneration in the private forests of Gadoli and Manda Khal, Pauri Garhwal, Uttarakhand, India.

**Figure 10.**
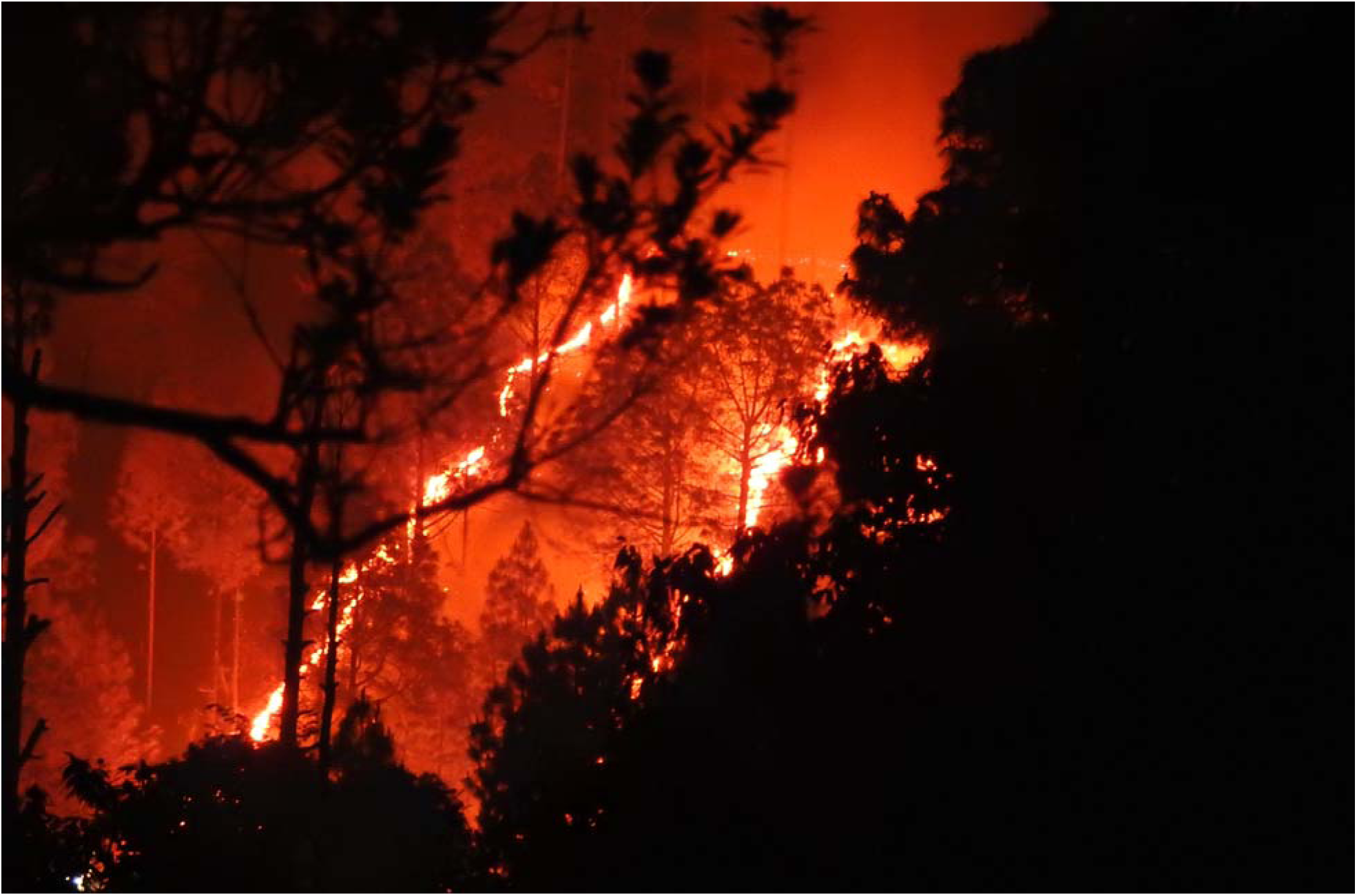
A forest fire on the periphery of the Gadoli and Manda Khal Fee Simple Estates, Pauri Garhwal, Uttarakhand, India

Confiscated weapons, such as *daranti’s* (traditional sickles) (Fig.11.) used to cut forest regrowth, highlight the threat to forest regeneration.

**Figure 11.**
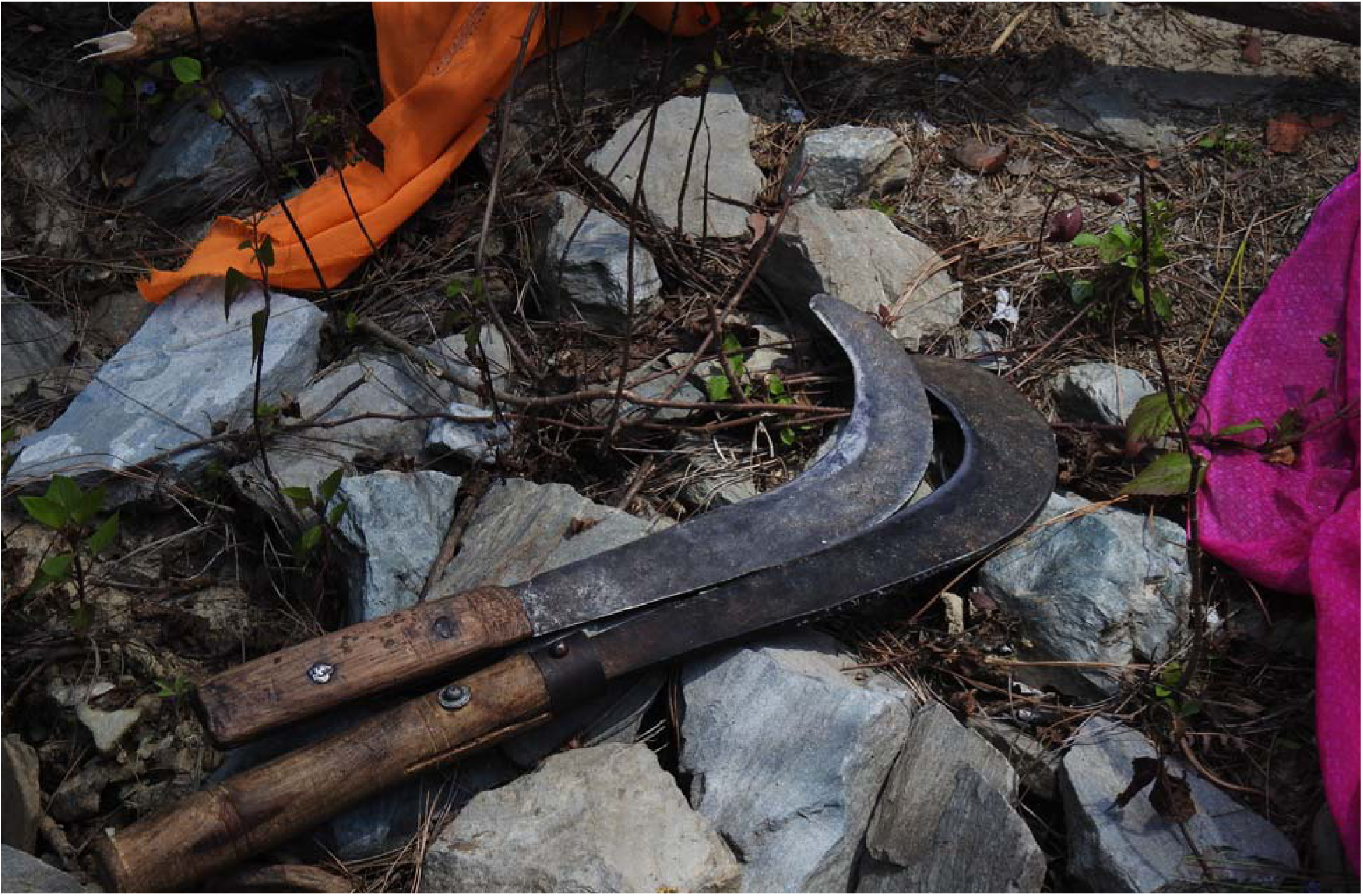
Confiscated weapons (locally known as Daranti’s) used for lopping and cutting forest regeneration.

Patrolling has significantly reduced illegal activities, improving habitat quality and extent [Chowfin & Leslie, 2021]. This has led to increased sightings of herbivores like barking deer (*Muntiacus muntjak*) (Fig. 12a & 12b) (Video File 5: https://doi.org/10.6084/m9.figshare.23620626.v1), pheasants (*Lophura leucomelanos*) (Fig. 13) (Video File 6: https://doi.org/10.6084/m9.figshare.23620626.v1) wild boar (*Sus scrofa*) (Fig 14), and Himalayan langur (*Semnopithecus schistaceus*) (Fig. 15.), which are considered prey species for leopards in this landscape.

**Figure 12a & 12b.**
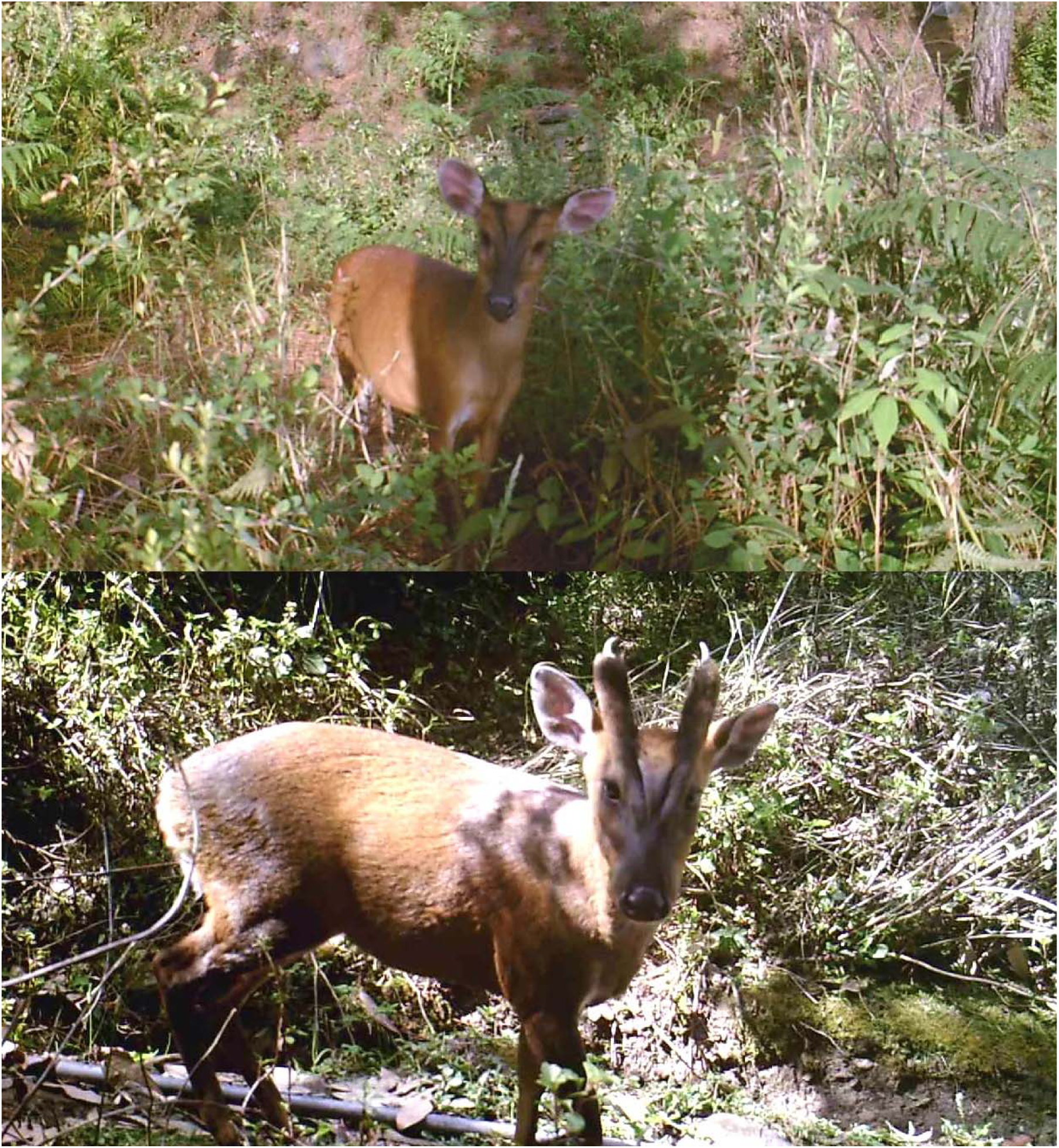
Barking deer (Muntiacus muntjak) in the private forests of the Gadoli and Manda Khal Fee Simple Estates, Pauri Garhwal, Uttarakhand, India.

**Figure 13.**
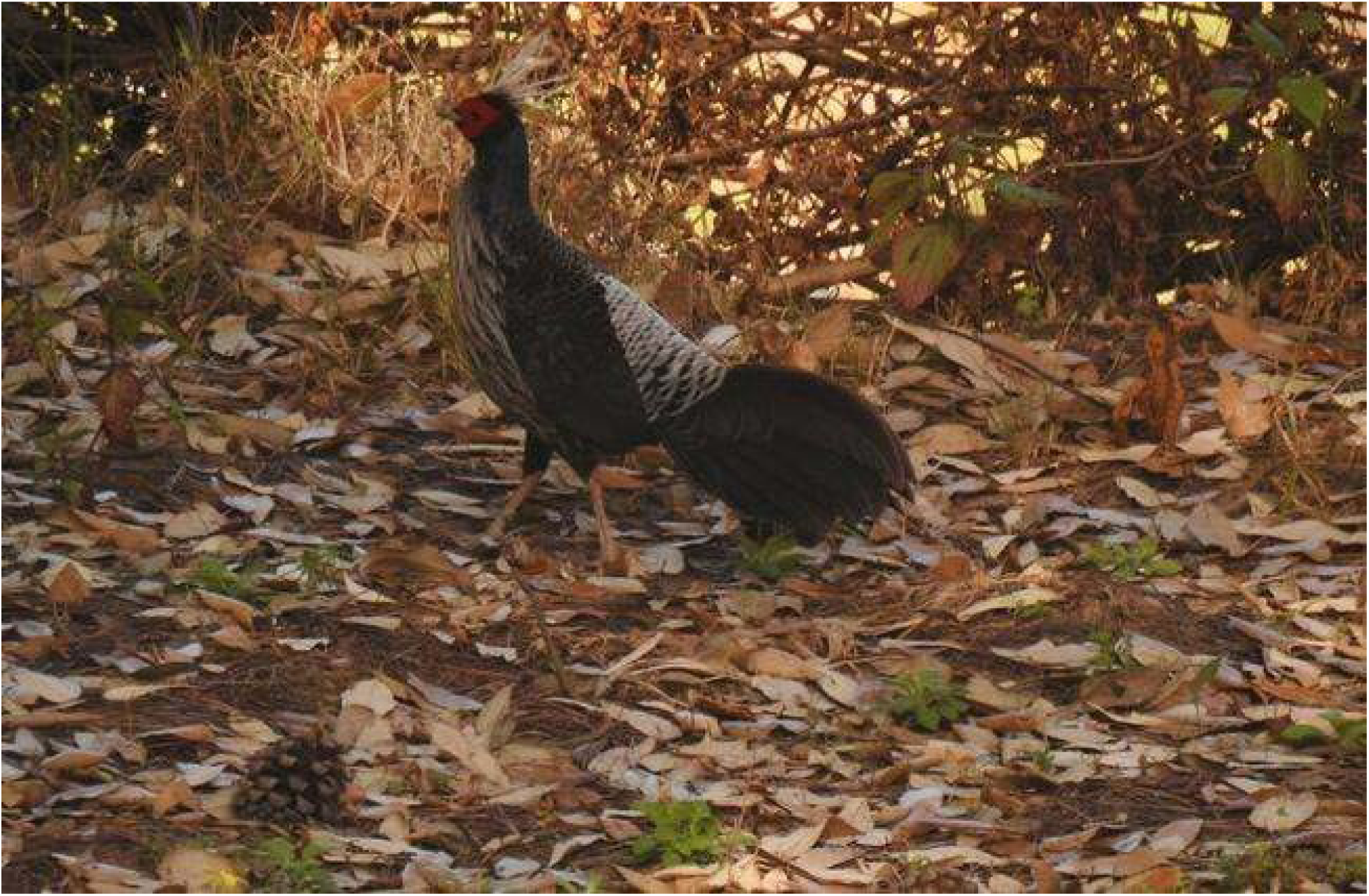
Khaleej pheasant (Lophura leucomelanos) in the private forests of the Gadoli and Manda Khal Fee Simple Estates, Pauri Garhwal, Uttarakhand, India.

**Figure 14.**
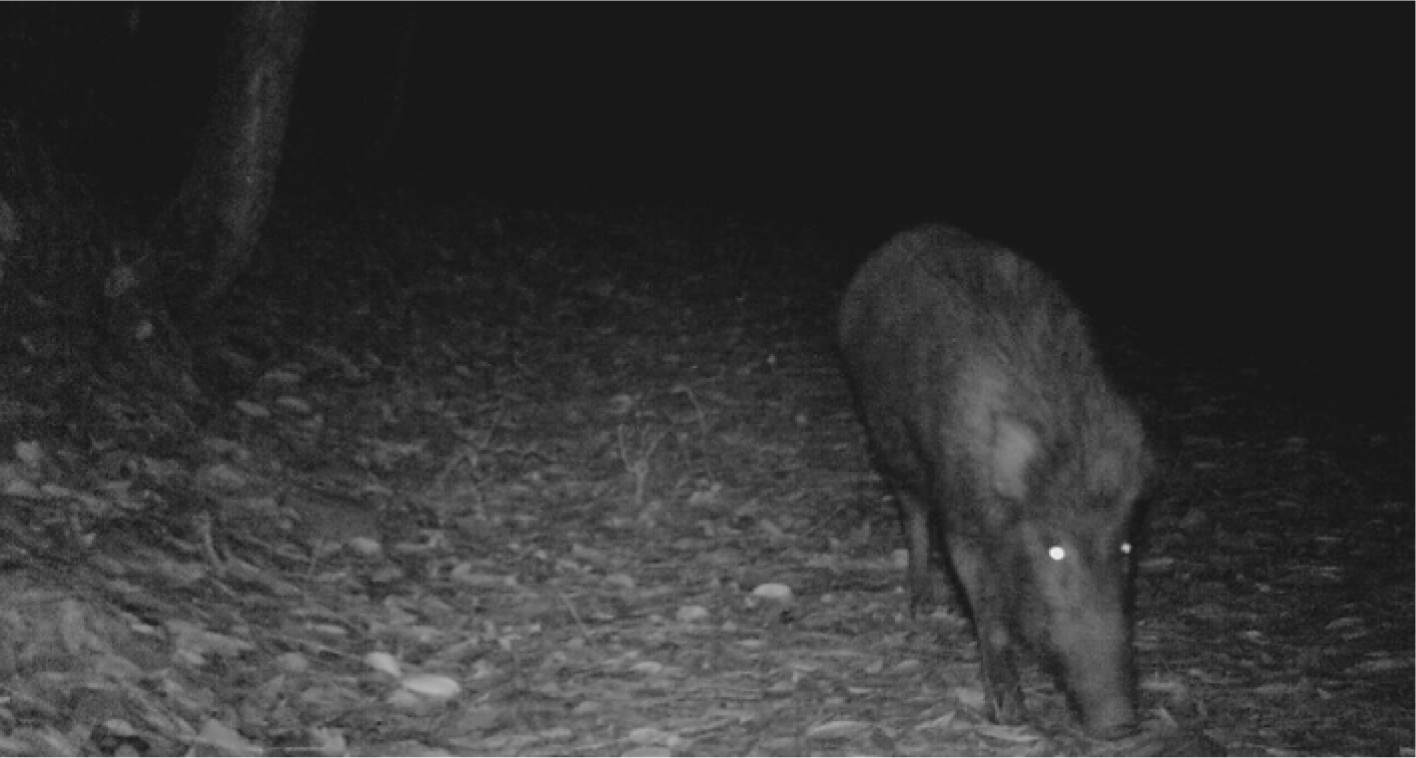
Wild Boar (Sus scorfa) in the private and Manda Khal Fee Simple Estates, Pauri Garhwal, Uttarakhand, India

**Figure 15.**
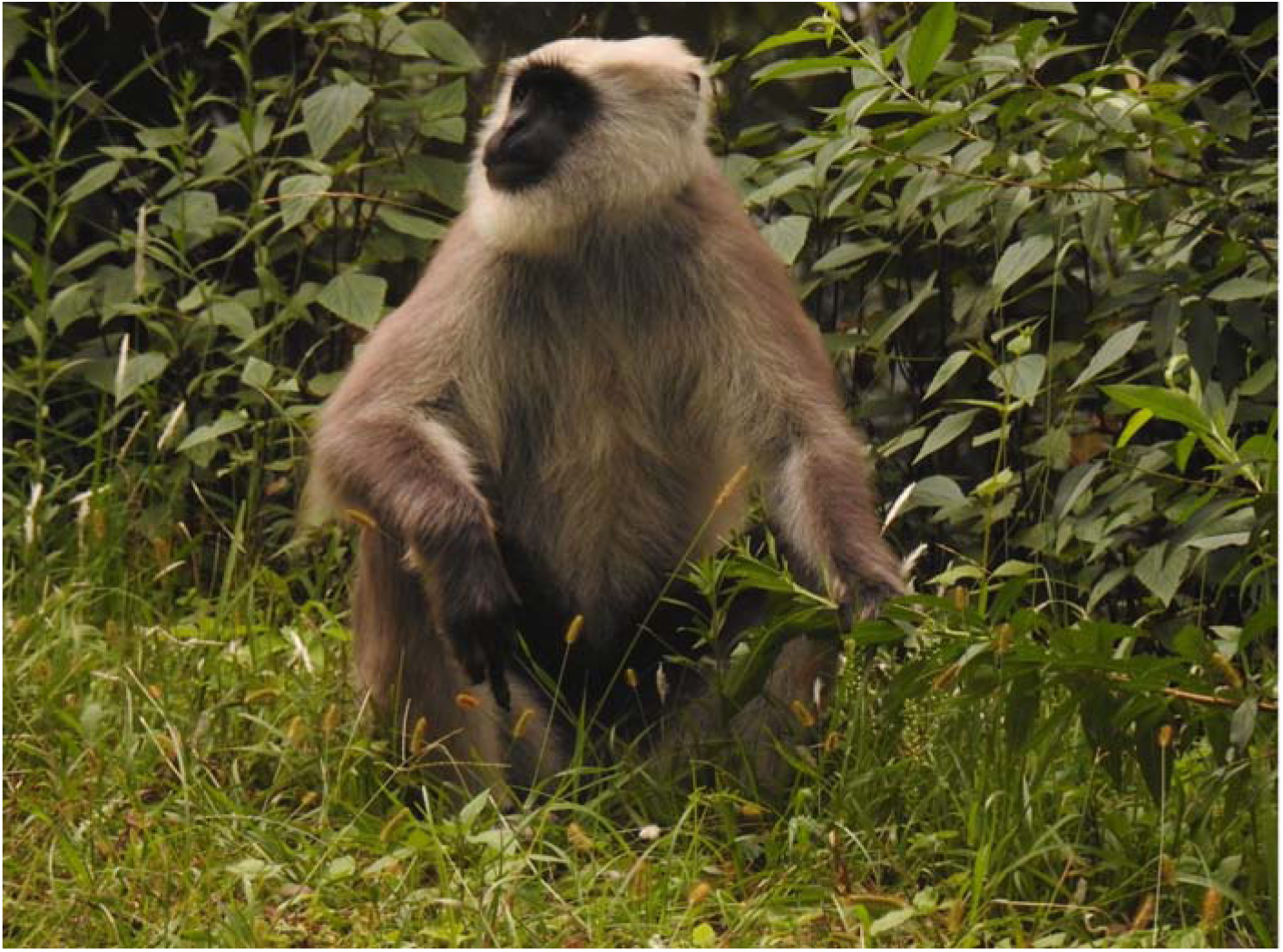
Himalayan Langur (Semnopithecus schistaceus) in the Gadoli and Manda Khal Fee Simple Estates, Pauri Garhwal, Uttarakhand, India.

### 4.3. Data compilation and analysis

For the purpose of this study, human - leopard conflict incidents were defined as leopard attacks on people resulting in the death or injury of the person attacked. If an attack resulted in the death of a person it was considered a fatality and if the attack resulted in an injury it was considered a non-fatal incident. Data on victims from leopard attacks were sourced from the Uttarakhand Forest Department under India’s Right to Information Act, 2005 [Ministry of Law and Justice, 2005]. This data included compensation records and forest range cases on fatal attacks and injuries to people by leopards for the Pauri Garhwal district. From this raw data, a district – level database (Supplementary Material A-i: https://doi.org/10.6084/m9.figshare.23620626.v1) was compiled spanning a 21-year period for the years 2000-2020 which consisted of both fatal and non-fatal incidents totaling 290 reported incidents in the Pauri Garhwal district over this time - period.

The district – level database was curated and tabulated into datasets and sub-datasets based on complete – case analysis (i.e. where incidents with missing data are excluded) [Everitt & Hothorn, 2011] to analyze human - leopard conflict incidents with inferences on its patterns according to the following categories:

i. Average patterns in human – leopard conflict with time-series analysis
ii. Gender - based patterns in human – leopard conflict with time – series analysis
iii. Habitat – based patterns in human – leopard conflict
iv. Age-based patterns in human – leopard conflict
v. Season - based patterns in human – leopard conflict with time-series analysis
vi. Daily temporal – based patterns in human – leopard conflict

Each category was curated into datasets and sub-datasets (Supplementary Material B: https://doi.org/10.6084/m9.figshare.23620626.v1) and groups were derived from sub-datasets for analysis.

To examine differences in human–leopard conflict incidents across categorical predictors such as gender, age group, habitat, season, and incident type, Generalized Linear Models (GLMs) were used, where, the distribution family (Poisson or Negative binomial) was selected based on the dispersion parameter.

To determine the dispersion parameter, initially, a Poisson GLM was fitted to the count data. The assumption of equidispersion (variance equal to the mean) was assessed based on the dispersion statistic, which is calculated as the ratio of the residual deviance to the degrees of freedom. If the dispersion parameter exceeded 1.5, which indicated overdispersion, a negative binomial GLM was used instead.

Model coefficients were interpreted on the log scale, and exponentiated values were reported as rate ratios with 95% confidence intervals. Pairwise comparisons between categorical groups were undertaken and model summaries included intercept and coefficient estimates with standard errors and p-values, and confidence intervals for exponentiated values. Where applicable, annual averages were also derived from model intercepts to aid interpretation.

When comparing groups, the null hypothesis (H_o_) assumed that there was no difference between the groups (p – value > 0.05, alpha = 5%) and the alternate hypothesis (p-value < 0.05, alpha = 5%) suggested that there was a significant difference between the groups.

Time-series analysis for annual averages, gender and season from 2000 – 2020 (Supplementary Material C: https://doi.org/10.6084/m9.figshare.23620626.v1) was performed using annual time – steps to determine if there was a statistically meaningful trend in the time – series based on r² and p-values using a Microsoft Excel 2007 add-in by Bryhn and Dimberg [2011]. In a time-series analysis, if one or several regressions concerning time and values in a time series, or time and mean values from intervals into which the series has been divided, yields r²≥0.65 and p≤0.05 then the time series is statistically meaningful [Bryhn & Dimberg, 2011] and indicates a trend in the time-series. Time-series analysis of human - leopard conflict incidents by habitat, age-group, time of attacks and their interactions with each other and with other categories were not performed due to missing data which could lead to biased inferences.

Human – leopard conflict incidents of the two most affected age – groups (i.e. children and adults) were studied to infer if belonging to a particular gender resulted in the type of incident (i.e. fatal or non – fatal) from a leopard attack. A binary logistic regression, which is a special case of a generalized linear model (GLM), was performed, from which predicted probabilities were derived. A threshold of 0.5 was set to classify predicted probabilities as belonging to the fatal or non – fatal category where a predicted probability of ≥ 0.5 indicated being in the non – fatal category and a predicted probability of < 0.5 indicated being in the fatal category. Human – leopard conflict incidents were studied to understand the influence of season on the gender of victims, incident type, habitat and age groups using binary or multinomial logistic regression depending on the type of response, from which predicted probabilities were derived and categorized into respective categories using a threshold of 0.5.

All statistical analysis were performed in the R Statistical Environment [R Core Team, 2023] in combination with packages “FSA” [Ogle et al., 2022] “rstatix” [Kassambara, 2022], “MASS” [Venables & Ripley, 2002], “nnet” [Venables & Ripley, 2002] and “emmeans” [Lenth, R. 2025]. For time in a 24 –hour period when leopard attacks took place, data was curated and analyzed in R with packages “astroFns” [Harris, 2012] and “activity” [Rowcliffe, 2021].

To ascertain if regulating the entry of people in forests in the district was reflected in any manner in human - leopard conflict, a comparison was made between the period from mid-April to mid-July 2020, which coincided with the three-month global lockdown due to the COVID-19 pandemic, and the data for the same months from 2000 to 2019. Specifically, the data for mid-April to mid-July from 2000 to 2019 was organized, tabulated and examined, and then juxtaposed with the incidents of human-leopard conflicts during the corresponding period in 2020.

Drivers of human – leopard conflict in the district were classified to provide insights on the causes of this conflict. Human - leopard conflict in the Pauri sub-division was compared with the Gadoli and Manda Khal Fee Simple Estates, located in the central himalayan tracts of the Pauri sub-division in the district of Pauri Garhwal, which is subject to ecological restoration – ecosystem recovery.

All Supplementary Information have been provided on figshare repository (https://doi.org/10.6084/m9.figshare.23620626.v1) which includes video files. For databases, readers may refer to Supplementary Material A. Detailed results with analysis’, datasets and sub – datasets are provided as Supplementary Material B. Data for time – series analysis are provided as Supplementary Material C.

## 5. Results

### 5.1 Average patterns in human – leopard conflict with time series analysis

A total of 290 human–leopard conflict incidents were analyzed between 2000 and 2020 representing 100% of all recorded incidents. During this period, the annual average of human– leopard conflict incidents (both fatalities and non-fatal incidents) across the district was 13.81 per year, with fluctuations over time. Time-series analysis showed no statistically significant trend between 2000 and 2020 (time – series: r² = 0.004, p = 0.775), indicating no evidence of an increase or decrease in human–leopard conflict over the two decades. Fatalities accounted for 22.41% (n=65/290) of incidents, with an annual average of 3.10 fatalities per year. Time-series analysis revealed no statistically significant trend for fatalities between 2000 and 2020 (time – series: r² = 0.021, p = 0.530), suggesting no increase or decrease in human – leopard conflict with respect to fatalities during this time - period. Non-fatal incidents comprised 77.59% (n=225/290) of conflicts, with an annual average of 10.71 incidents per year. Similarly, time-series analysis for non-fatal incidents showed no significant trend (time – series: r² = <0.001, p = 0.982), indicating no increase or decrease in human – leopard conflict with respect to non – fatal incidents with respect to this time – period. Generalized linear models (GLMs) were used to analyze the effect of incident type (fatal vs. non-fatal) on the annual number of leopard attack victims (Table 3). The results indicated a statistically significant difference in victims between the two incident types (β = 1.242, SE = 0.194, p < 0.001). The estimated rate ratio (RR) for non-fatal relative to fatal incidents was RR = 3.46 (95% CI: 2.38–5.08), demonstrating that non-fatal attacks occurred more than three times as often as fatal ones. The model intercept (β = 1.130, SE = 0.156, p < 0.001) represents the expected number of fatal incidents per year and was significantly greater than zero, suggesting a stable average of approximately 3.1 fatal attacks annually. These findings confirm that while both types of incidents persisted over the years, non-fatal encounters were substantially more frequent than fatal ones and that human–leopard conflict was significantly skewed toward non-fatal incidents.

**Table 3.**
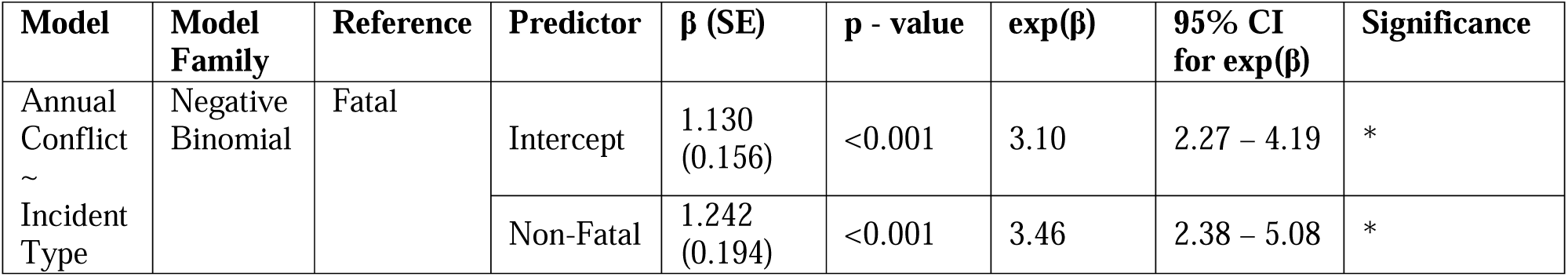
Annual human – leopard conflict (GLM)

### 5.2 Gender – based patterns in human – leopard conflict with time-series analysis

A total of 290 human–leopard conflict incidents were analyzed by gender between 2000 and 2020. 53% were male (n = 153) and 47% were female victims (n = 137). Generalized linear models (GLMs) were used to evaluate the effects of gender on outcome (fatal vs. non-fatal). There was no statistically significant difference in the overall number of incidents between male and female victims or when modeling the type of incident by gender, indicating that conflict incidence was evenly distributed across genders. However, significant differences were observed between fatal and non – fatal incidents (female: p = 0.0029; male: p = <0.001) for both genders during this time period (Table 4).

**Table 4.** Gender – based patterns in human – leopard conflict (GLMs)

| Model | Model Family | Reference | Predictor | $\beta$ (SE) | p - value | exp($\beta$) | 95% CI for exp($\beta$) | Significance |
| --- | --- | --- | --- | --- | --- | --- | --- | --- |
| Conflict ~ Gender | Negative Binomial | Female | Intercept | 1.876 (0.129) | <0.001 | 6.52 | 5.07 – 8.41 | * |
|  |  |  | Male | 0.111 (0.180) | 0.540 | 1.12 | 0.78 – 1.59 | NS |
| Female Victims ~ Incident Type | Negative Binomial | Fatal | Intercept | 0.644 (0.223) | 0.0038 | 1.90 | 1.37 – 2.56 | * |
|  |  |  | Non-Fatal | 0.865 (0.291) | 0.0029 | 2.38 | 1.65 – 3.47 | * |
| Male Victims ~ Incident Type | Negative Binomial | Fatal | Intercept | 0.174 (0.214) | 0.414 | 1.19 | 0.76 – 1.77 | NS |
|  |  |  | Non-Fatal | 1.633 (0.243) | <0.001 | 5.12 | 3.23 – 8.40 | * |
| Fatal ~ Gender | Negative Binomial | Female | Intercept | 0.644 (0.265) | 0.015 | 1.90 | 1.15 – 3.25 | * |
|  |  |  | Male | -0.470 (0.394) | 0.233 | 0.63 | 0.29 – 1.35 | NS |
| Non-Fatal ~ Gender | Negative Binomial | Female | Intercept | 1.509 (0.137) | <0.001 | 4.52 | 3.45 – 5.91 | * |
|  |  |  | Male | 0.314 (0.186) | 0.092 | 1.37 | 0.95 – 1.97 | NS |

Time-series analysis indicated no statistically significant trend for either gender across this two-decade period (males: r² = 0.011, p = 0.647; females: r² < 0.001, p = 0.962).

### 5.3 Habitat – based patterns in human – leopard conflict

Spatial patterns of human - leopard conflict were studied by categorizing incidents by habitat. A total of 250 incidents (86% of the complete dataset) were included in the habitat analysis, with 42% (n = 106) accounted for in forests and 58% (n = 144) in human habitation.

Generalized linear models (GLMs) were used to assess the influence of habitat type on leopard conflict frequency. The results indicated no statistically significant difference between habitat types (β = 0.306, SE = 0.192, p = 0.111). The rate ratio (RR) for human habitation to forest was RR = 1.36 (95% CI: 0.93–1.98), which indicated a 36% higher frequency of incidents in human habitation when compared to forests, but this difference was not statistically significant. The model intercept (β = 1.619, SE = 0.140, p < 0.001) reflects an average of approximately 5.05 conflict incidents per year in forest habitats. Statistically significant differences were observed in fatalities (irrespective of gender) (p = <0.001), female fatalities (p < 0.001), male fatalities (p = 0.003) and male non – fatal incidents (p = 0.035) between human habitation and forest (Table 5).

**Table 5.**
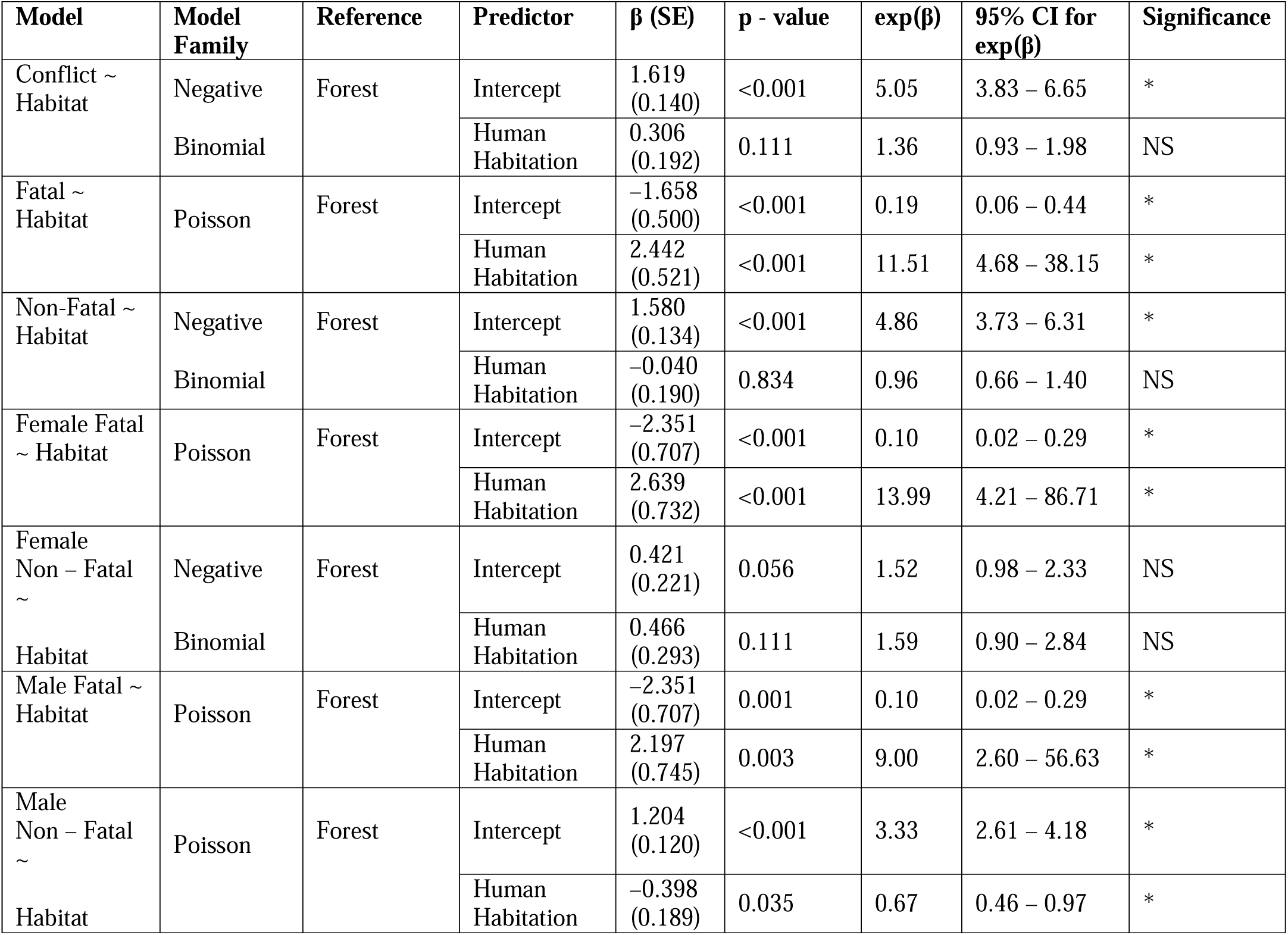
Habitat – based patterns in human – leopard conflict (GLMs)

### 5.4 Age-based patterns in human – leopard conflict

Human – leopard conflict incidents were categorized into age-groups to understand patterns in human – leopard conflict incidents ordered by age – group and consisted of 275 incidents representing 95% (n=275/290) of the total incidents from 2000-2020. Age-groups were categorized into children (1-12 years), teen (13-19), adult (20-60) and aged (61 and above).

Children accounted for 29% (n=79/275), teens 10% (n=28/275), adults 55% (n=150/275), and the aged 6% (n=18/275) of incidents. Generalized linear models (GLMs) were used to assess the effect of age group (child, teen, aged vs. adult) on the number of human–leopard conflict incidents (Table 6).

**Table 6.** Age-based patterns in human – leopard conflict (GLMs)

| Model | Model Family | Reference | Predictor | $\beta$ (SE) | p - value | exp($\beta$) | 95% CI for exp($\beta$) | Significance |
| --- | --- | --- | --- | --- | --- | --- | --- | --- |
| Conflict ~<br>Age Group | Negative Binomial | Adult | Intercept | 1.897<br>(0.158) | <0.001 | 6.67 | 4.92 – 9.16 | * |
|  |  |  | Aged | –1.946<br>(0.305) | <0.001 | 0.14 | 0.08 – 0.26 | * |
|  |  |  | Child | –0.572<br>(0.236) | 0.015 | 0.56 | 0.35 – 0.90 | * |
|  |  |  | Teen | –1.684<br>(0.285) | <0.001 | 0.19 | 0.10 – 0.32 | * |

The results indicated that incidents differed significantly by age group. Compared to adults, incidents involving aged victims were significantly lower (β = –1.946, SE = 0.305, p < 0.001). The estimated rate ratio (RR) for aged individuals relative to adults was RR = 0.14 (95% CI: 0.08–0.26), indicating that conflict incidents were substantially less frequent among aged victims. Similarly, conflict incidents were significantly lower for teens compared to adults (β = −1.684, SE = 0.285, p < 0.001; rate ratio = 0.19, 95% CI: 0.10–0.32), and also significantly lower for children (β = –0.572, SE = 0.236, p = 0.015; rate ratio = 0.56, 95% CI: 0.35–0.90). The model intercept (β = 1.897, SE = 0.158, p < 0.001) represents the expected number of adult victims annually, at approximately 6.67 incidents annually, indicating that adults experienced the highest number of leopard conflict incidents, These findings indicate that adults were disproportionately affected compared to other age groups.

Overall, human–leopard conflict incidents were significantly skewed towards adults and children, with adults being the most affected group, followed by children.

For children, a total of 79 incidents (n = 79) were categorized by gender and incident type. Male children accounted for 41% (n=32/79) of incidents, while female children accounted for 59% (n=47/79). Among male victims, 40.63% (n=13/32) were fatalities, and 59.37% (n=19/32) were non-fatal incidents. Among female victims, 51.06% (n=24/47) were fatalities, and 48.93% (n=23/47) were non-fatal incidents.

A binary logistic regression was conducted on the child group, to assess the effect of gender on the type of incident (fatal vs. non-fatal) during leopard attacks, with incident type as the response variable and gender as the predictor. The equation derived was:

*Y(Non-Fatal) = 0.3795 – 0.4220 * female*

Predicted probabilities indicated that for male children, the likelihood of a non-fatal incident was 0.5938, suggesting non-fatal incidents were more likely than fatalities. For female children, the probability was 0.4939, indicating fatalities were more likely. The negative coefficient for females shows that the odds of a non-fatal incident decrease when the victim is female compared to males.

Among the 150 adult incidents (n = 150), 62% involved males (n = 93/150), while 38% involved females (n = 57/150). Among male victims, 5.38% (n = 5/93) were fatalities, and 94.62% (n = 88/93) were non-fatal incidents. Among female victims, 10.53% (n = 6/57) were fatalities, and 89.47% (n = 51/57) were non-fatal incidents.

A binary logistic regression was conducted on adults, to examine gender in relation to the type of incident (fatal vs. non-fatal), with incident type as the response variable and gender as the predictor. The derived equation was:

*Y(Non-Fatal) = 2.8679 – 0.7278 * female*

The predicted probability of a non-fatal incident was 0.9462 for males, indicating non-fatal incidents were more likely than fatalities for adult males. For females, the predicted probability was 0.8947, also indicating non-fatal incidents were more likely. The negative coefficient suggests that the odds of a non-fatal incident decrease for females compared to males.

### 5.5 Season – based patterns in human – leopard conflict with time-series analysis

Human–leopard conflict incidents were analysed across seasons for all 290 tabulated incidents (100%). Seasons were categorized into Summer (March to mid-June), Monsoon (mid-June to September), and Winter (October to February). Of all incidents, 28% occurred in summer (n=81), 30% in monsoon (n=88), and 42% in winter (n=121). The seasonal distribution of incidents followed the pattern: summer < monsoon < winter. Generalized Linear Models (GLMs) were used to interpret the effect of season on different categories (Table 7).

**Table 7.** Season – based patterns in human – leopard conflict (GLMs)

| Model | Model Family | Reference | Predictor | $\beta$ (SE) | p - value | exp($\beta$) | 95% CI for exp($\beta$) | Significance |
| --- | --- | --- | --- | --- | --- | --- | --- | --- |
| Conflict ~ Season | Negative Binomial | Summer | Intercept | 1.350<br>(0.145) | <0.001 | 3.85 | 2.89 – 5.12 | * |
|  |  |  | Monsoon | 0.083<br>(0.203) | 0.683 | 1.09 | 0.73 – 1.62 | NS |
|  |  |  | Winter | 0.401<br>(0.195) | 0.040 | 1.49 | 1.02 – 2.19 | NS |
| Female ~ Season | Negative Binomial | Summer | Intercept | 0.452<br>(0.207) | 0.029 | 1.57 | 1.03 – 2.34 | * |
|  |  |  | Monsoon | 0.167<br>(0.285) | 0.558 | 1.18 | 0.68 – 2.07 | NS |
|  |  |  | Winter | 0.678<br>(0.266) | 0.011 | 1.97 | 1.17 – 3.34 | * |
| Male ~ Season | Negative Binomial | Summer | Intercept | 0.827<br>(0.175) | <0.001 | 2.29 | 1.61 – 3.20 | * |
|  |  |  | Monsoon | -0.087<br>(0.251) | 0.729 | 0.92 | 0.56 – 1.50 | NS |
|  |  |  | Winter | 0.189<br>(0.240) | 0.431 | 1.21 | 0.76 – 1.94 | NS |
| Fatal ~ Season | Poisson | Summer | Intercept | 0.000<br>(0.209) | 0.663 | 1.00 | 0.71 – 1.61 | NS |
|  |  |  | Monsoon | -0.191<br>(0.310) | 0.538 | 0.83 | 0.45 – 1.51 | NS |
|  |  |  | Winter | -0.000<br>(0.295) | 1.000 | 1.00 | 0.56 – 1.79 | NS |
| Non – Fatal ~ Season | Negative Binomial | Summer | Intercept | 1.050<br>(0.160) | <0.001 | 2.86 | 2.08 – 3.89 | * |
|  |  |  | Monsoon | 0.140<br>(0.222) | 0.528 | 1.15 | 0.75 – 1.78 | NS |
|  |  |  | Winter | 0.470<br>(0.212) | 0.027 | 1.60 | 1.06 – 2.43 | * |
| Female Fatal ~ Season | Negative Binomial | Summer | Intercept | -0.647<br>(0.380) | 0.089 | 0.52 | 0.24 – 1.09 | NS |
|  |  |  | Monsoon | 0.310<br>(0.514) | 0.547 | 1.36 | 0.50 – 3.80 | NS |
|  |  |  | Winter | 0.241<br>(0.519) | 0.642 | 1.27 | 0.46 – 3.57 | NS |
| Female Non – Fatal ~ Season | Negative Binomial | Summer | Intercept | 0.000<br>(0.245) | 1.000 | 1.00 | 0.60 – 1.58 | NS |
|  |  |  | Monsoon | 0.251<br>(0.330) | 0.447 | 1.29 | 0.67 – 2.48 | NS |
|  |  |  | Winter | 0.847<br>(0.304) | 0.005 | 2.33 | 1.30 – 4.30 | NS |
| Male Fatal ~ Season | Poisson | Summer | Intercept | -0.847<br>(0.333) | 0.011 | 0.43 | 0.21 – 0.77 | * |
|  |  |  | Monsoon | -0.588<br>(0.558) | 0.292 | 0.56 | 0.17 – 1.61 | NS |
|  |  |  | Winter | 0.201<br>(0.450) | 0.655 | 1.22 | 0.51 – 3.03 | NS |
| Male Non – Fatal ~ Season | Poisson | Summer | Intercept | 0.619<br>(0.160) | <0.001 | 1.86 | 1.33 – 2.50 | * |
|  |  |  | Monsoon | 0.074<br>(0.222) | 0.739 | 1.08 | 0.70 – 1.67 | NS |
|  |  |  | Winter | 0.187<br>(0.217) | 0.389 | 1.21 | 0.79 – 1.85 | NS |

In terms of annual averages, the summer season recorded 3.86 incidents per year (time – series: r² = 0.007, p = 0.726), the monsoon season recorded 4.19 incidents per year (time – series: r² = 0.002, p = 0.840), and the winter season recorded 5.76 incidents per year (time – series: r² < 0.001, p = 0.933). Time-series analysis indicated that conflict remained constant across all seasons over the two-decade period. GLMs revealed a significant seasonal effect, with winter emerging as a period of elevated conflict. Compared to summer, monsoon incidents were not significantly different (β = 0.083, SE = 0.203, p = 0.683), with a rate ratio (RR) of RR = 1.09 (95% CI: 0.73–1.62). Winter incidents were significantly higher (β = 0.401, SE = 0.195, p = 0.040), RR = 1.49 (95% CI: 1.02–2.19). A comparison between monsoon and winter revealed no significant difference (β = 0.318, SE = 0.192, p = 0.098), RR = 1.37 (95% CI: 0.89–2.19). The model intercept (β = 1.350, SE = 0.145, p < 0.001) also indicated an estimated ∼3.86 incidents per year in summer.

A binary logistic regression was conducted to assess the influence of seasons on the gender of leopard attack victims, with gender (male vs. female) as the response variable and season as the predictor (Gender ∼ Season). The equation derived was:

*Y(Female) = −0.2955 + 0.2955 * monsoon + 0.4008 * winter*

The predicted probabilities indicated that in summer, females were less likely to be victims (0.4267), in monsoon, both genders had equal likelihood (0.5), and in winter, females were more likely to be victims (0.5263).The analysis suggests that the probability of female victims increased from summer to winter, with higher odds in winter compared to summer and monsoon. A binary logistic regression was conducted to assess the influence of seasons on the type of leopard attack incident (fatal vs non-fatal), with incident type as the response variable and season as the predictor (Incident Type ∼ Season). The equation derived was:

*Y(Non-Fatal) = 1.0809 + 0.3054 * monsoon + 0.5159 * winter*

The predicted probabilities, for summer (0.7467), monsoon (0.80) and winter (0.8316) showed that non-fatal incidents were more likely than fatalities in all seasons. The analysis suggests that the likelihood of non-fatal incidents increased from summer to winter, with higher odds in the winter compared to summer and the monsoon.

A binary logistic regression was conducted to assess the influence of seasons on leopard attack locations (forest vs. human habitation), with habitat as the response variable and season as the predictor (Habitat ∼ Season). The equation derived was:

*Y(Human Habitation) = 0.5754 - 0.1699 * monsoon - 0.6807 * winter*

The predicted probabilities indicated that leopard attacks were more likely to occur in human habitation during summer (0.64) and monsoon (0.60), whereas in winter (0.4737), attacks were more likely to occur in forests. The analysis revealed that leopard attacks in human habitation decreased from summer to winter, while attacks in forests increased. Negative coefficients for monsoon and winter indicated lower odds of attacks in human habitation compared to summer.

A multinomial logistic regression was performed to understand the manner in which seasons had an influence on age - groups. Keeping age - group as the response variable and season as the predictor variable (Age Group ∼ Season), the multinomial logistic regression equations derived were:

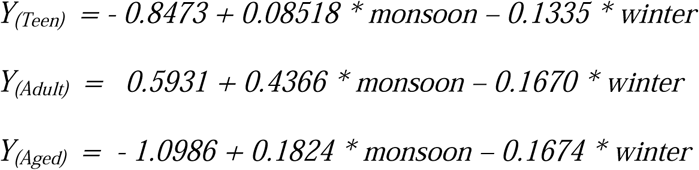

from which predicted probabilities were derived (Table 8).

**Table 8.** Summary of predicted probabilities.

| **Season** | **P(Child)** | **P(Teen)** | **P(Adult)** | **P(Aged)** |
| --- | --- | --- | --- | --- |
| Summer | 0.2802 | 0.1201 | 0.5064 | 0.0933 |
| Monsoon | 0.2142 | 0.1 | 0.6 | 0.0858 |
| Winter | 0.3136 | 0.1177 | 0.4805 | 0.0882 |

For children, the highest predicted probability occurs in the winter. For teens, the predicted probabilities of being victims of leopard attacks indicate that teens were less likely to be affected than children in all seasons, with the highest predicted probabilities being in summer and winter. A positive monsoon coefficient suggests higher odds of teen victims compared to summer. However, due to the combined effects of other coefficients and the baseline category, this does not translate into a higher predicted probability. The negative winter coefficient reflects lower odds of teens being victims in winter compared to summer, but the predicted probability for teens in winter is similar to summer due to the interplay of probabilities among all groups.

For adults, the predicted probabilities indicate that adults were more likely to be victims than children across all seasons, with the highest predicted probabilities being in the monsoon and summer. The positive monsoon coefficient reflects increased odds of adult victims compared to summer, consistent with the rise in predicted probability The negative winter coefficient, reflects decreased odds of adult victims in winter compared to summer, consistent with the lower predicted probability.

For the aged, the predicted probabilities indicate that children were more likely to be affected than the aged in all seasons, with low probabilities across all seasons. A positive monsoon coefficient shows increased odds of aged victims compared to summer. However, the predicted probability does not reflect this increase because of the relative contributions of all groups. The negative winter coefficient suggests lower odds in winter compared to summer, consistent with the predicted probability.

### 5.6 Daily temporal – based patterns in human – leopard conflict

To understand patterns in incidents in a 24-hour period, data on the time when leopard attacks occurred were tabulated and consisted of 233 incidents for representing 80.34% of the total number of incidents from 2000 – 2020. Data which was in hh:mm was converted into radians with the R package “astroFns” and then analyzed with the R package “activity”.

Attacks were concentrated between sunrise and sunset without discriminating between habitat and seasons with incidents peaking thrice in a 24-hour period. The first during in the morning (10:30 hours ± 30 minutes), the second in the afternoon (14:00 hours ± 30 minutes) and the third in the evening (18:00 hours ± 30 minutes) (Fig. 16)

**Figure 16.**
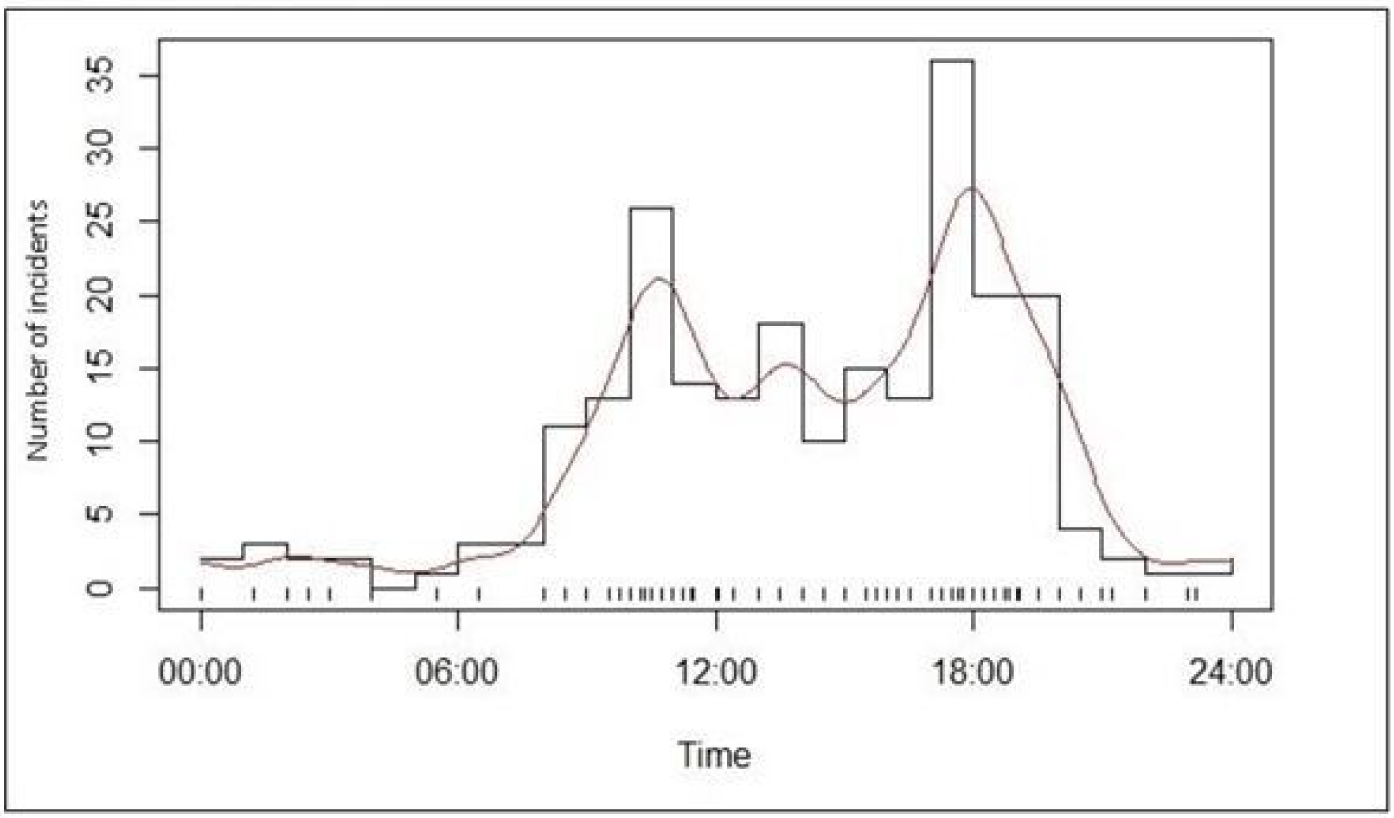
Overall(N = 233)

To determine the time when incidents occurred in forests without discriminating between seasons, incidents (n=91) were analyzed after removal of missing data. Incidents largely occurred between between 9.00 and 18.30 hours with a large number of incidents in between 10.00 and 18.00 hours (Fig. 17).

**Figure 17.**
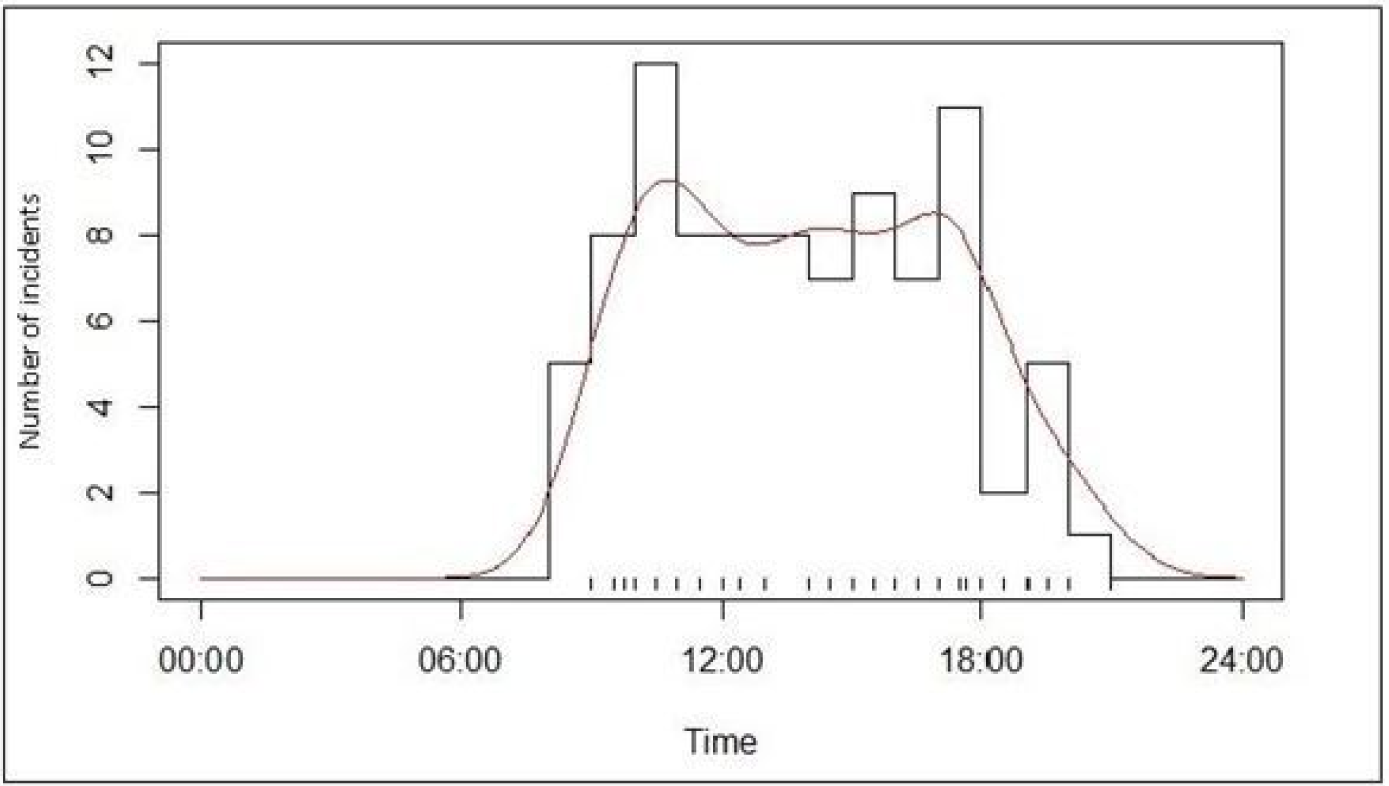
Forest(N = 91)

To determine time when incidents occurred in human habitation without discriminating between seasons, incidents (n = 132) were analyzed after removal of missing data. Incidents largely occurred between 6:00 hours and 22:00 hours peaking thrice during this period. The first during in the morning (10:30 hours ± 30 minutes), the second in the afternoon (14:00 hours ± 30 minutes) and the third in the evening (18:00 hours ± 30 minutes) (Fig. 18).

**Figure 18.**
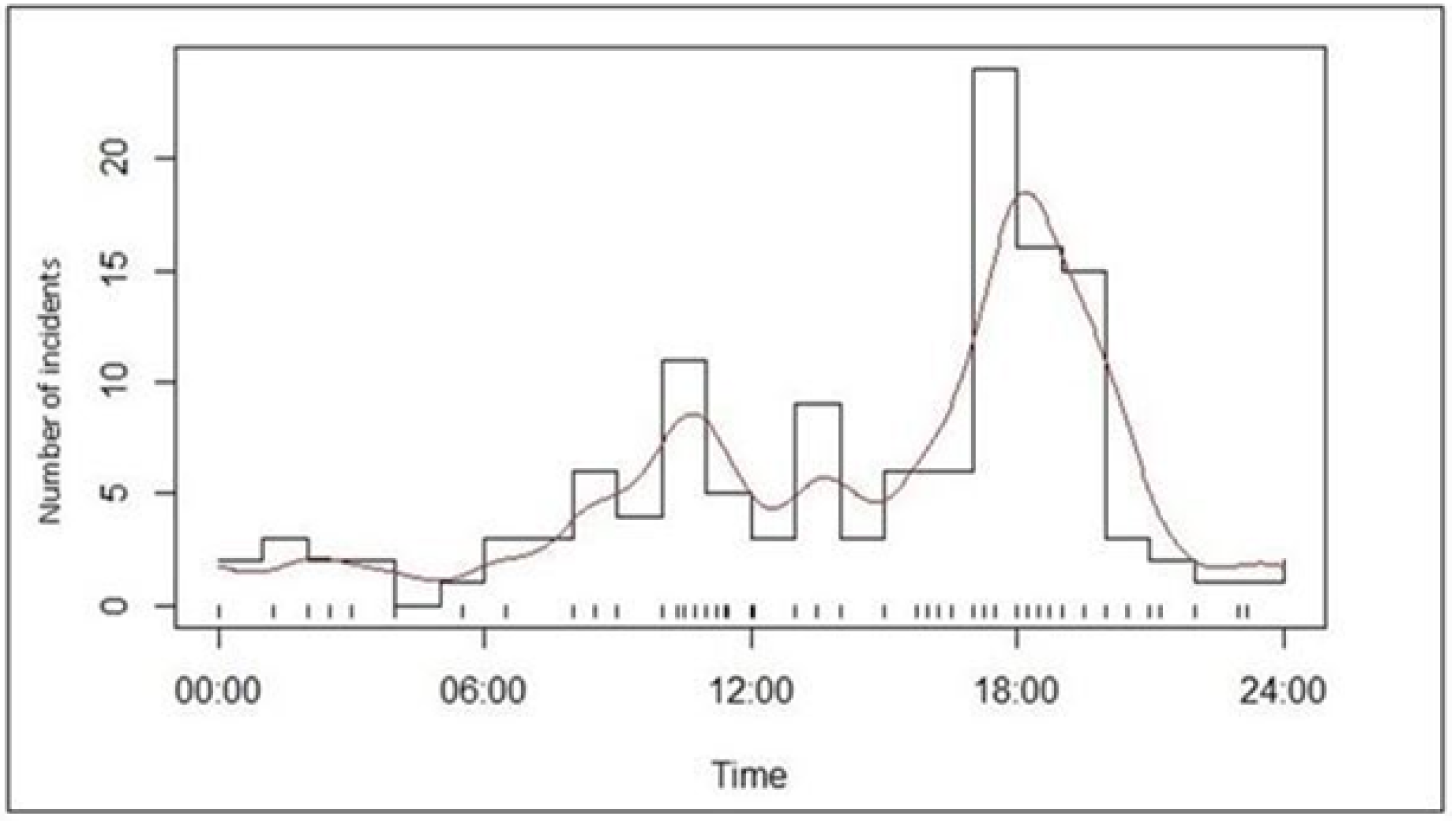
Human Hbbitation (N = 132)

Incident times were analyzed by season without differentiating between habitat. For summer (n = 66), incidents primarily occurred between 7:00 and 21:00 hours, peaking twice: first around 11:00 hours and then again at approximately 19:30 hours, with several incidents in between (Fig. 19).

**Figure 19.**
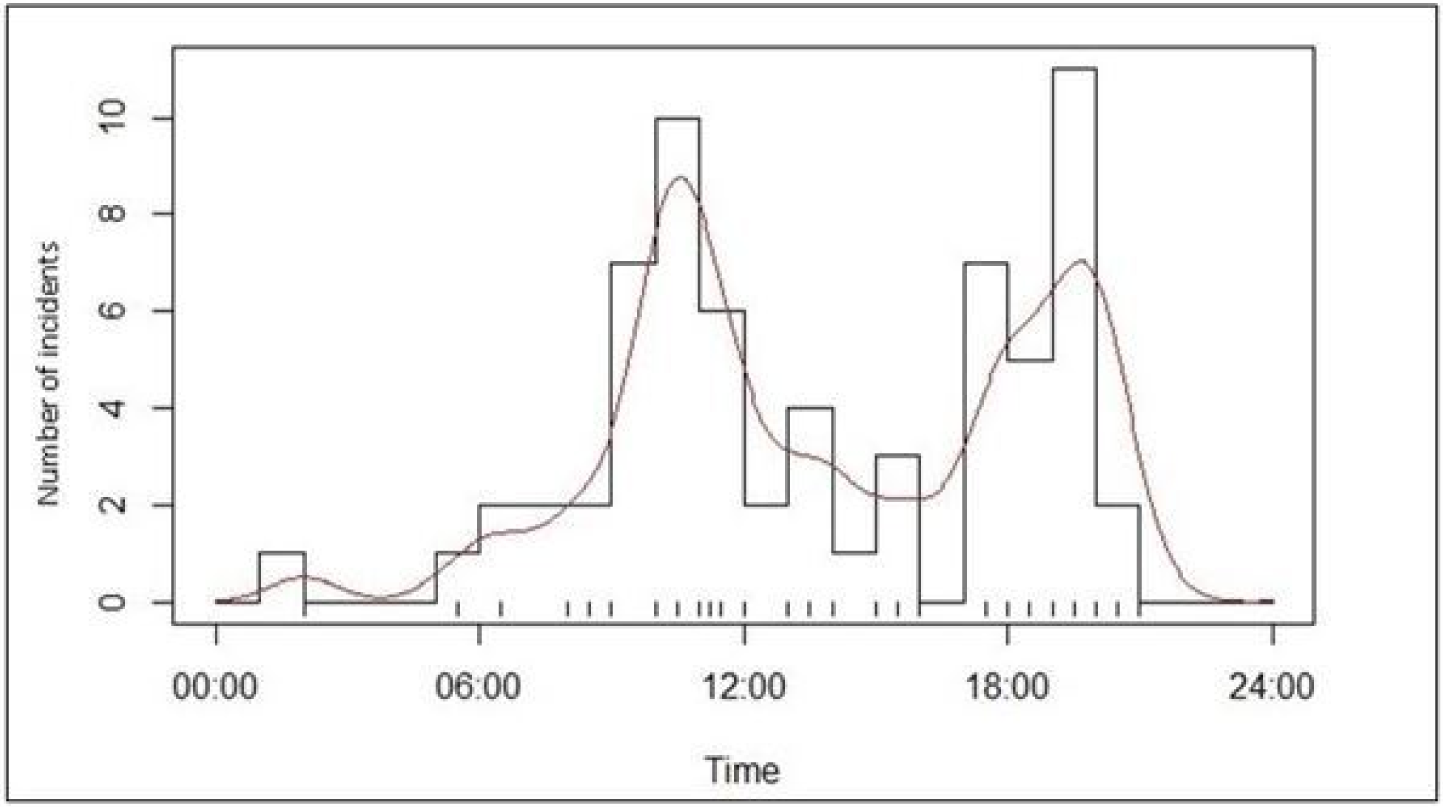
Summer (N = 66)

For monsoon (n = 75), incidents occurred largely between 9:00 and 19:00 hours, peaking at 11:00 hours and 18:30 hours, with other incidents scattered throughout this period (Fig.20).

**Figure 20.**
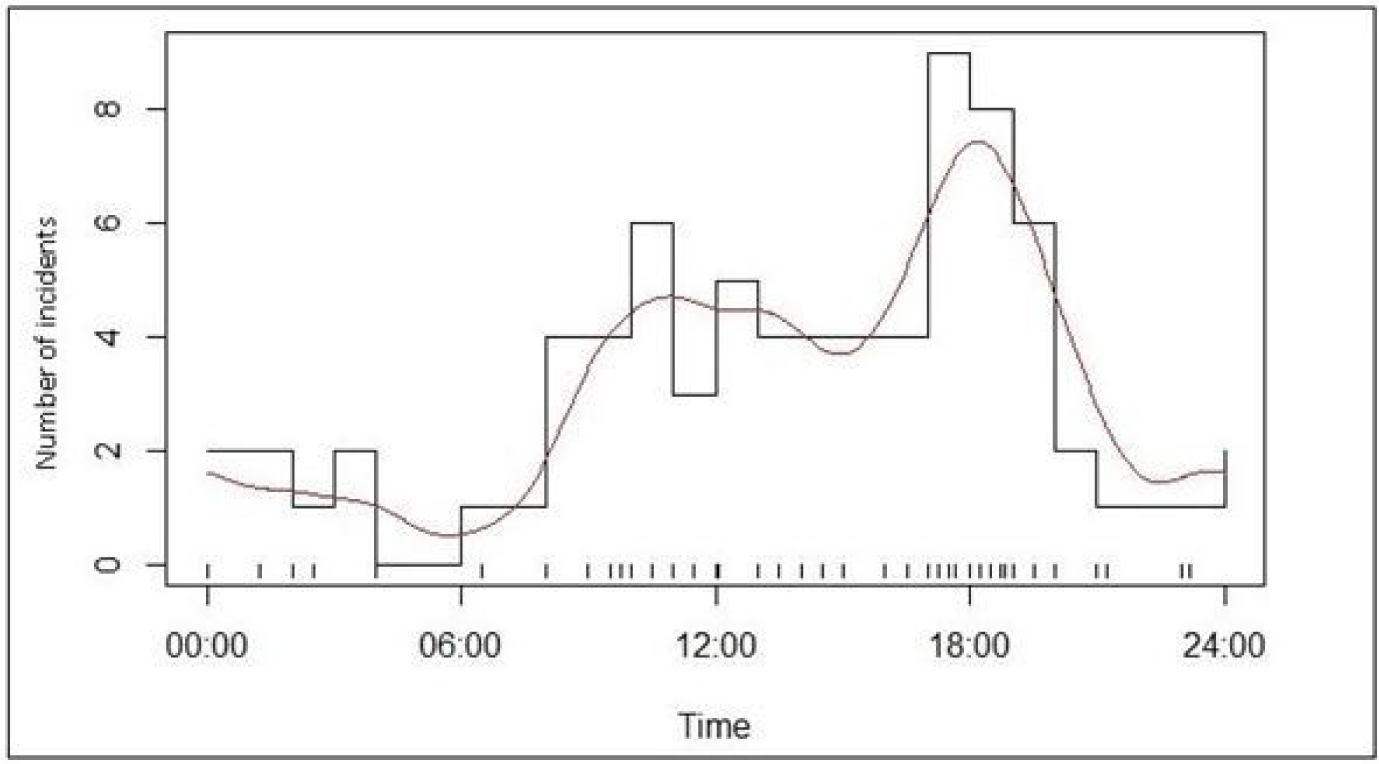
Monsoon(N = 75)

For winter (n = 92), incidents increased between 9:00 and 18:30 hours, with a peak at 18:00 hours (Fig.21).

**Figure 21.**
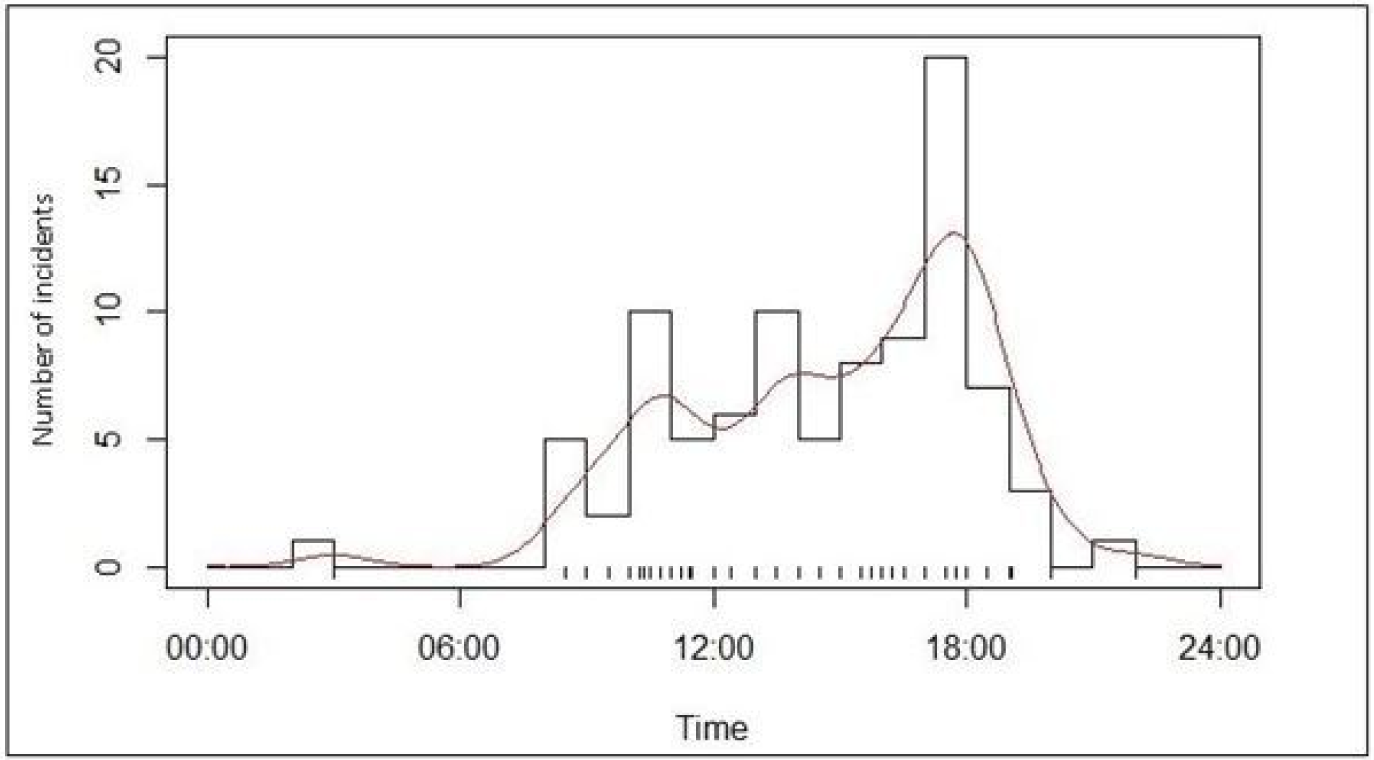
Winter(N = 92)

Incident times were analyzed by habitat and season. In human habitation, during summer (n = 42), incidents primarily occurred between 06:00 and 20:00 hours, with two peaks: around 11:00 hours and 18:30 hours (Fig.22).

**Figure 22.**
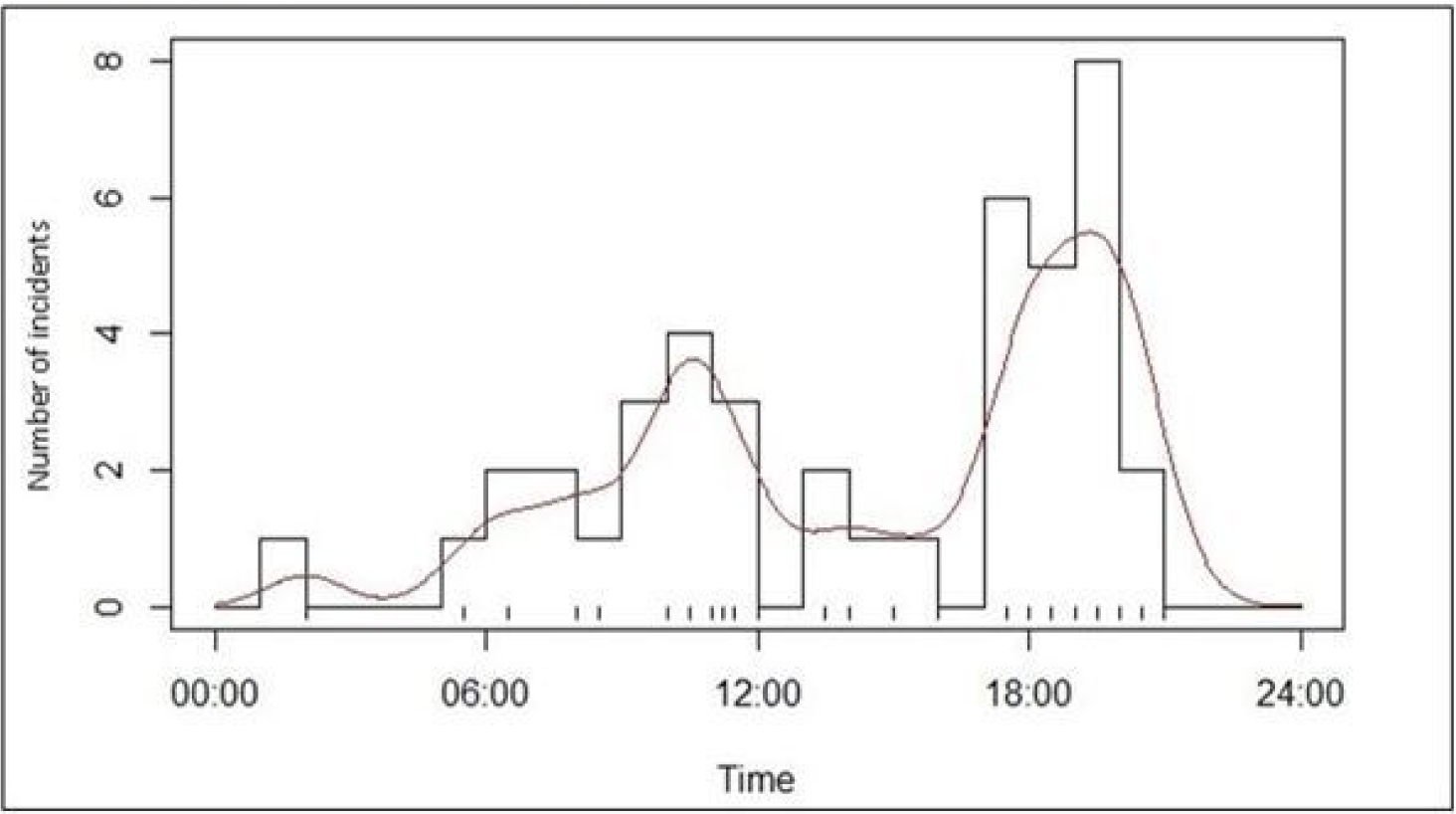
Human Habitation – Summer(N= 42)

In the monsoon (n = 45), incidents were spread throughout the 24-hour period, with peaks at 13:00 hours and 19:30 hours (Fig.23).

**Figure 23.**
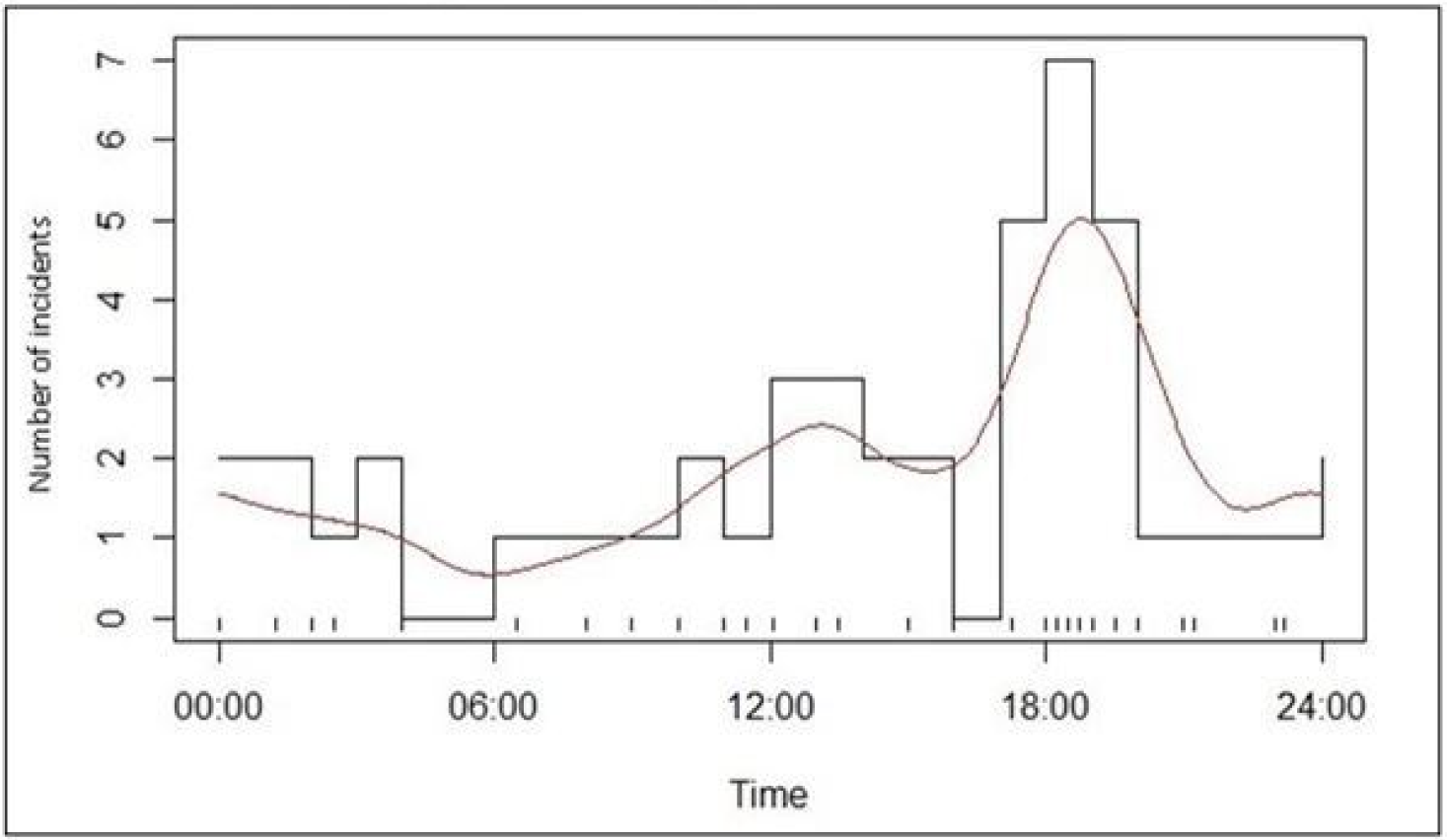
Human Habitation – Monsoon(N=45)

In winter (n = 43), incidents occurred mainly between 08:00 and 23:00 hours, peaking at approximately 10:30 hours and 18:00 hours (Fig. 24).

**Figure 24.**
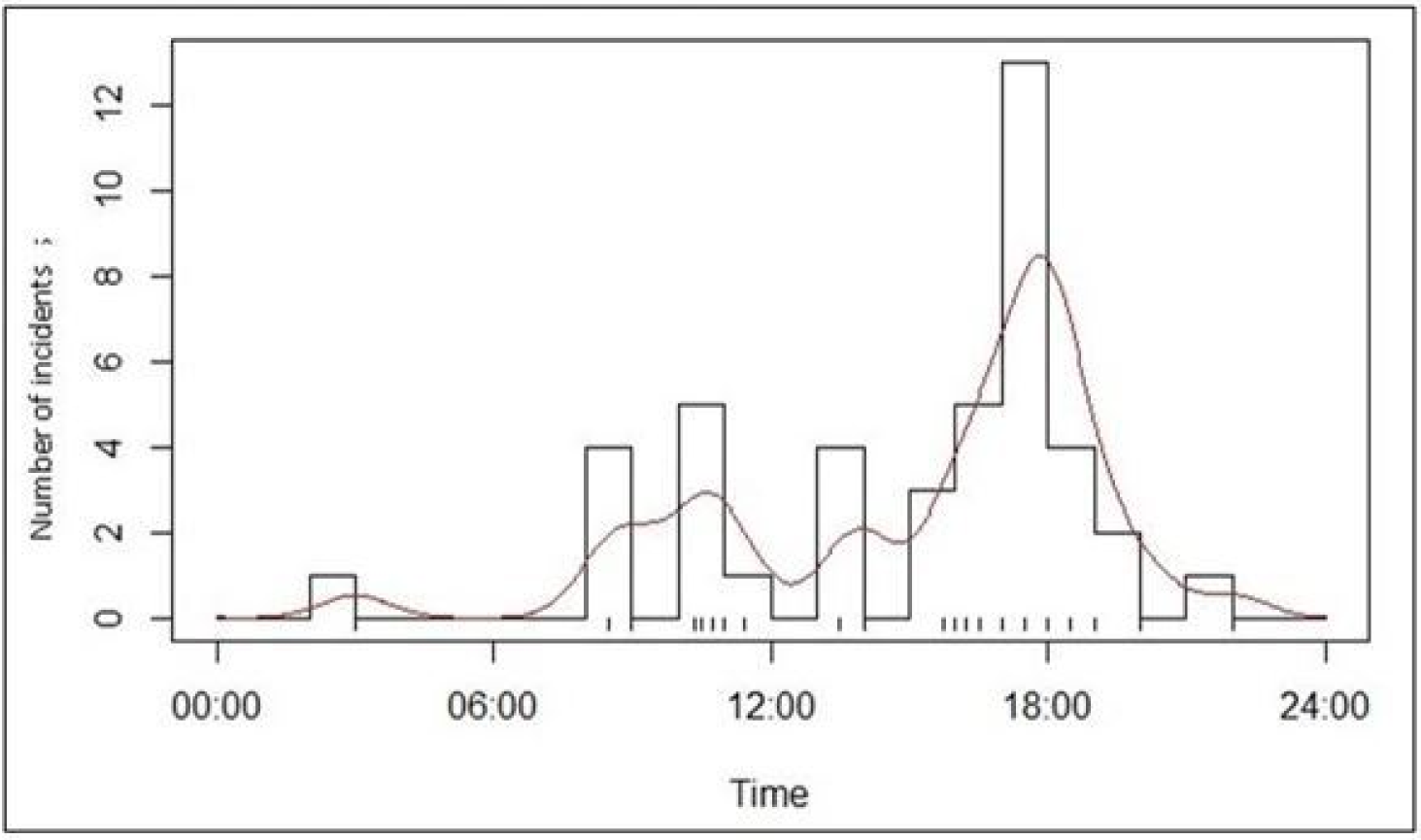
Human Habitation – Winter(N=43)

In forests, during summer (n = 23), incidents mainly occurred between 9:00 and 20:00 hours, with two peaks: around 11:30 hours, followed by a decline, and again at 19:30 hours (Fig. 25).

**Figure 25.**
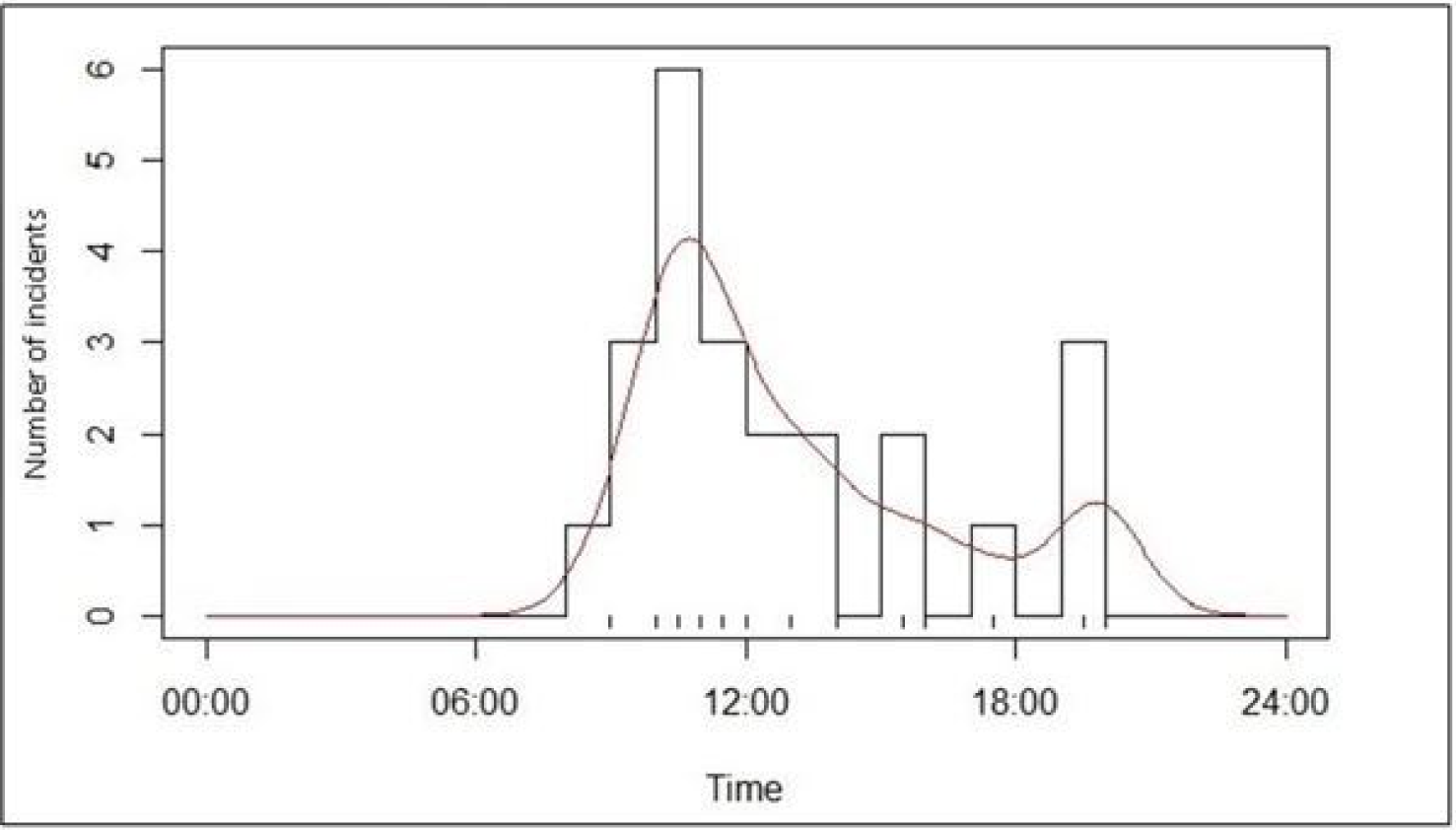
Forest – Summer(N=23)

In the monsoon (n = 26), incidents occurred mostly between 9:00 and 21:00 hours, peaking at 10:30 hours, declining, and then peaking again at 17:00 hours (Fig.26).

**Figure 26.**
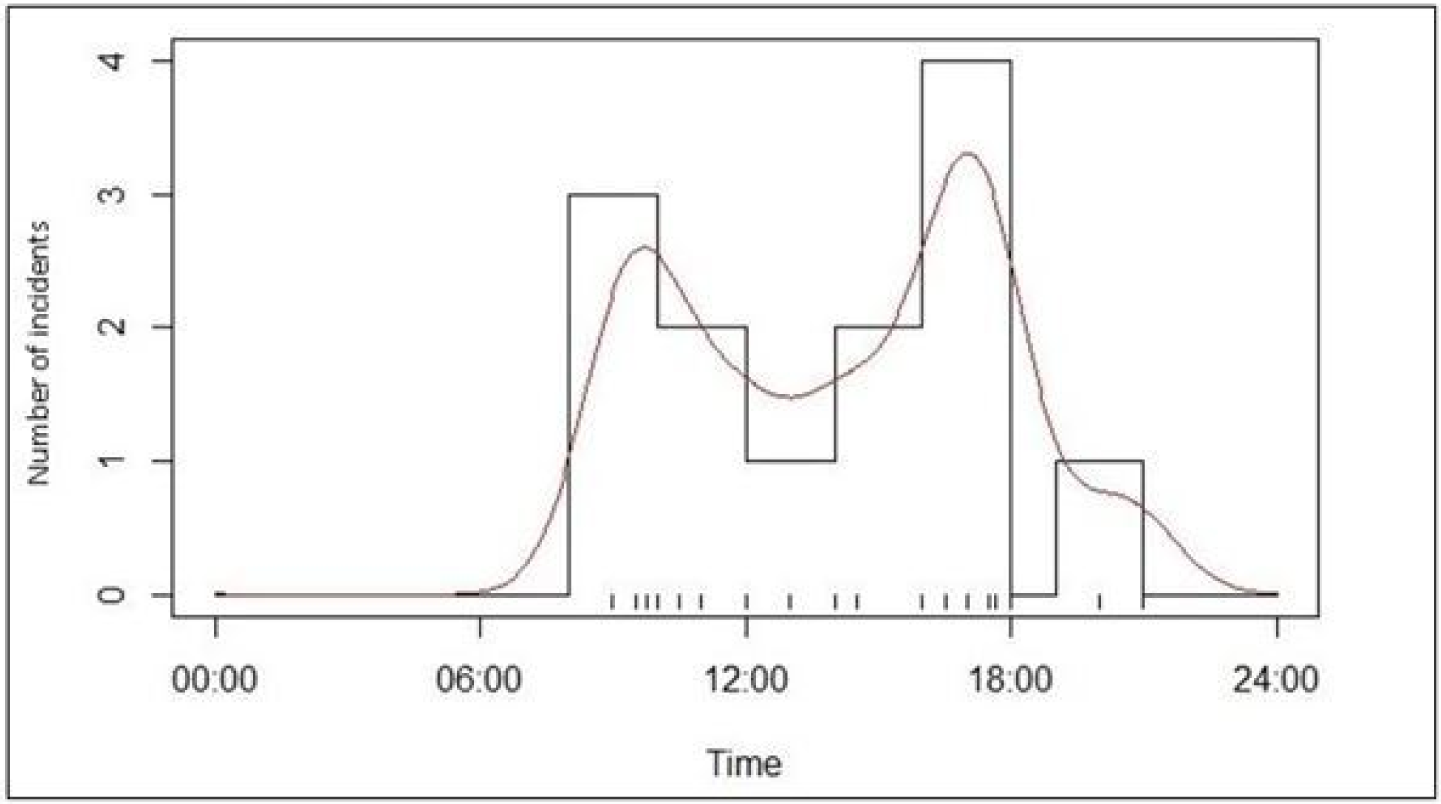
Forest – Monsoon(N=26)

In winter (n = 42), incidents occurred predominantly between 9:00 and 19:00 hours, with a peak between 14:00 and 17:30 hours (Fig.27).

**Figure 27.**
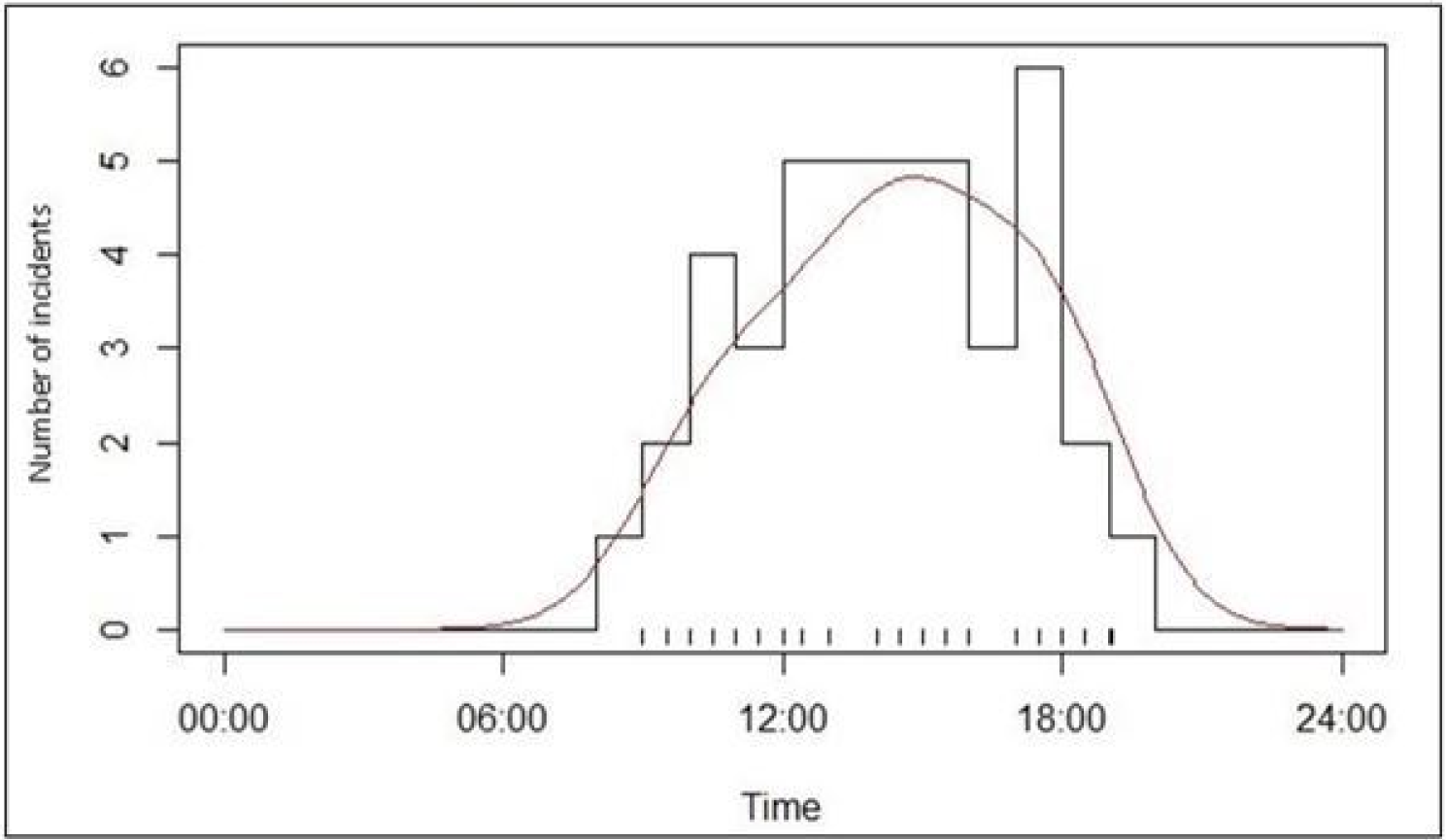
Forest – Winter(N = 42)

### 5.7 Human – leopard conflict in the Pauri Sub – Division and the Gadoli and Manda Khal Fee Simple Estates

The Pauri sub-division (Fig. 28) with a geographical extent of 1594.64 square kilometers represents 29.92% (n=1594.64/5329) of the geographical area of the Pauri Garhwal district.

**Figure 28.**
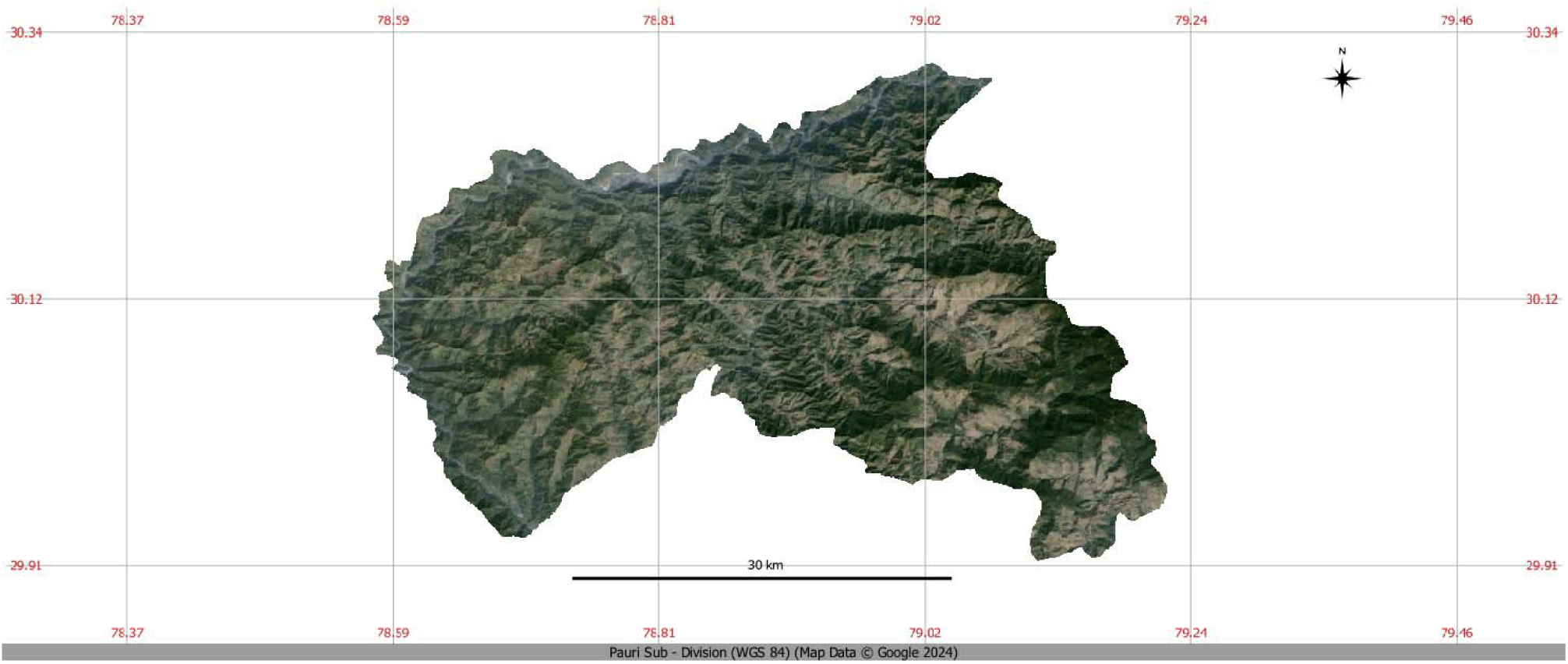
Map of the Pauri sub – division in the Pauri Garhwal District, Uttarakhand, India

The total population in the sub-division is estimated to be 135,718 persons with 72,140 females and 63,578 males at a sex ratio (per 1000 persons) of 1135 females per 1000 males and a population density of 177 persons / km^2^ [DEIAA, 2018]. Of the 290 human – leopard conflict incidents tabulated for the district of Pauri Garhwal, 232 incidents occurred in the Pauri sub-division accounting for 80% (n=232/290) of human – leopard conflict incidents in the district (Supplementary Material A – ii: https://doi.org/10.6084/m9.figshare.23620626.v1). From 2000-2020, the annual average of human-leopard conflict incidents in the Pauri sub-division was 11.05, with no significant trend (time – series: r²=0.024, p=0.499). Fatalities totaled 48 incidents (74% of district fatalities; n=48/65), averaging 2.29 per year, with no significant trend (time – series: r²=0.011, p=0.652). Non-fatal incidents numbered 184 (82% of district non-fatal incidents; n = 184/225), averaging 8.76 per year, with no significant trend (time – series: r²=0.021, p=0.528). In contrast, the Gadoli and Manda Khal Fee Simple Estates, located at the conflict epicenter, reported no human-leopard conflict incidents over the same period. (Table 9).

**Table 9.** Comparison of human - leopard conflict between the Gadoli and Manda Khal Fee Simple Estates, the Pauri Sub-division and the Pauri Garhwal District (2000 – 2020)

| Year | Gadoli and Manda Khal Fee Simple Estates |  |  | Pauri Sub-division |  |  | Pauri Garhwal District |  |  |
| --- | --- | --- | --- | --- | --- | --- | --- | --- | --- |
|  | Fatal | Non-Fatal | Total | Fatal | Non-Fatal | Total | Fatal | Non-Fatal | Total |
| 2000 | 0 | 0 | 0 | 0 | 1 | 1 | 0 | 2 | 2 |
| 2001 | 0 | 0 | 0 | 2 | 6 | 8 | 5 | 11 | 16 |
| 2002 | 0 | 0 | 0 | 4 | 16 | 20 | 7 | 16 | 23 |
| 2003 | 0 | 0 | 0 | 3 | 5 | 8 | 4 | 9 | 13 |
| 2004 | 0 | 0 | 0 | 1 | 4 | 5 | 1 | 4 | 5 |
| 2005 | 0 | 0 | 0 | 1 | 16 | 17 | 1 | 18 | 19 |
| 2006 | 0 | 0 | 0 | 6 | 12 | 18 | 7 | 12 | 19 |
| 2007 | 0 | 0 | 0 | 4 | 9 | 13 | 5 | 9 | 14 |
| 2008 | 0 | 0 | 0 | 1 | 10 | 11 | 1 | 10 | 11 |
| 2009 | 0 | 0 | 0 | 1 | 15 | 16 | 1 | 16 | 17 |
| 2010 | 0 | 0 | 0 | 2 | 7 | 9 | 2 | 7 | 9 |
| 2011 | 0 | 0 | 0 | 1 | 16 | 17 | 1 | 16 | 17 |
| 2012 | 0 | 0 | 0 | 4 | 13 | 17 | 4 | 14 | 18 |
| 2013 | 0 | 0 | 0 | 6 | 6 | 12 | 6 | 8 | 14 |
| 2014 | 0 | 0 | 0 | 4 | 10 | 14 | 8 | 15 | 23 |
| 2015 | 0 | 0 | 0 | 4 | 15 | 19 | 7 | 18 | 25 |
| 2016 | 0 | 0 | 0 | 1 | 10 | 11 | 1 | 11 | 12 |
| 2017 | 0 | 0 | 0 | 0 | 2 | 2 | 0 | 6 | 6 |
| 2018 | 0 | 0 | 0 | 0 | 2 | 2 | 0 | 7 | 7 |
| 2019 | 0 | 0 | 0 | 3 | 6 | 9 | 3 | 10 | 13 |
| 2020 | 0 | 0 | 0 | 0 | 3 | 3 | 1 | 6 | 7 |
| **Total** | **0** | **0** | **0** | **48** | **184** | **232** | **65** | **225** | **290** |
| **Annual Average** | **0** | **0** | **0** | **2.29** | **8.76** | **11.05** | **3.10** | **10.71** | **13.81** |

### 5.8 Human – leopard conflict in the Pauri Garhwal district and the Coronavirus lockdown

During the 3-month global lockdown in 2020 (April-July), Pauri Garhwal enforced curfews implemented by the local police which restricted access to forests, effectively making them in-violate. Comparing human-leopard conflict data from mid-April to mid-July for 2000-2019, with that of 2020 for the same period (64 incidents, 22% of total incidents, average 3.20 per year; n = 64/290) no such incidents occurred in 2020 during the lockdown (Table 10).

**Table 10.** Comparison of human - leopard conflict between April to July (2000 – 2019) and April to July (2020)

| Year | Gadoli and Manda Khal Fee Simple Estates [April to July 2000-2019] | Pauri Garhwal District [April to July 2000 - 2019] |
| --- | --- | --- |
| 2000 | 0 | 0 |
| 2001 | 0 | 2 |
| 2002 | 0 | 7 |
| 2003 | 0 | 3 |
| 2004 | 0 | 2 |
| 2005 | 0 | 2 |
| 2006 | 0 | 5 |
| 2007 | 0 | 4 |
| 2008 | 0 | 3 |
| 2009 | 0 | 4 |
| 2010 | 0 | 1 |
| 2011 | 0 | 3 |
| 2012 | 0 | 6 |
| 2013 | 0 | 1 |
| 2014 | 0 | 1 |
| 2015 | 0 | 6 |
| 2016 | 0 | 3 |
| 2017 | 0 | 1 |
| 2018 | 0 | 3 |
| 2019 | 0 | 7 |
| **Total** | **0** | **64** |
| **Average** | **0** | **3.2** |
|  | Gadoli and Manda Khal Fee Simple Estates [April to July 2020] | Pauri Garhwal District [April to July 2020] |
| **Total** | **0** | **0** |
| **Average** | **0** | **0** |

However, after restrictions were lifted in mid-July 2020, human-leopard conflict resumed with incidents corresponding to the re-entry of people into forests.

## 6. Discussion

### 6.1. Key Findings and Novel Insights

#### 6.1.1. Human – Leopard Conflict History and Shifting Patterns

Human-leopard conflict in Garhwal has historical roots, dating back to the 1920s during the global influenza epidemic. The epidemic devastated the isolated Garhwal Himalayan region, predominantly inhabited by Hindus. With thousands of deaths, a shortage of wood for cremations led to human corpses being left along the Alaknanda River, which attracted leopards. One leopard developed a taste for human flesh by scavenging on these corpses, initiating attacks that lasted nine years until it was killed by Jim Corbett, as detailed in his book *The Man-Eating Leopard of Rudraprayag* [Corbett, 1948]. Corbett portrayed the leopard as a majestic creature, advocating for its conservation rather than animosity. Today, human-leopard conflicts occur on a larger scale, with heightened animosity toward leopards. Like the Rudraprayag leopard, many are killed by bullets or bait.

#### 6.1.2. Geographic and Socioeconomic Factors

Higher human population density in the Pauri sub-district compared to the rest of Pauri Garhwal has intensified forest degradation and deforestation, resulting in most conflicts occurring there. The presence of leopards near settlements suggests they are seeking prey in these areas. Expanding human settlements nearer to forests and forest encroachments further increase conflict risk.

#### 6.1.3. Conflict Distribution and Statistical Insights

The Pauri sub-division accounted for 80% (n=232/290) of human-leopard conflict incidents in the Pauri Garhwal district from 2000–2020, despite comprising only 29.92% of the district’s land area. However, in contrast, the Gadoli and Manda Khal Fee Simple Estates reported zero such incidents during this period. This absence of conflict, when compared to both the Pauri sub-division and on the larger scale the Pauri Garhwal district is highly significant both statistically and for conservation efforts. It however should be noted that on 23/10/2020, a single case of a teenage girl injured in a leopard attack was reported in the Manda Khal Fee Simple Estate but was excluded from the analysis. The incident occurred when the girl and a relative entered the forest to retrieve their cattle, which were grazing there illegally. As the leopard attacked the herd, the girl attempted to drive it away to protect the cattle and was subsequently injured. The nature of this attack appeared to be a defensive reaction by the leopard to provocation rather than a predatory attempt.

### 6.2. Demographic Vulnerability

#### 6.2.1. Demographics of Victims

From 1988–2000, human-leopard conflict in Pauri Garhwal disproportionately affected females (66%) and children under 15 years old (68%) [Chauhan et al., 2000; Ogra, 2008]. However, data from 2000–2020 reveals a shift: among 290 cases, 53% of victims were male (n=153) and 47% female (n=137). Adults were the most affected group, with males constituting 62% (n=93) of adult victims compared to 38% females (n=57). This male-biased pattern contrasts with earlier findings by Chauhan [2000] and Ogra [2008], despite the district’s female-biased human population.

#### 6.2.2. Age and Gender – Specific Vulnerabilities

Adults were the most frequently affected age group, with men experiencing more incidents than women, particularly in forested areas. This gender disparity may be attributed to men often traveling alone on forest paths for activities such as work, cattle grazing, or commuting between villages, thereby increasing their risk of encounters with leopards. In contrast, women typically enter forests in small groups for tasks like collecting fuelwood and fodder, which likely reduces their vulnerability to leopard attacks.

The majority of leopard attacks on children occurred within human habitation, potentially due to children being left unattended. Female children comprised 58% of the victims (n=46/79), compared to 42% male victims (n=33/79). Although no evidence links these incidents to the practice of female feticide or infanticide in Uttarakhand [Maini, 2019], the higher proportion of female victims raises significant concerns. The elevated fatality rates among children could be associated with parental negligence, as unattended children—particularly girls—may lack the physical ability to defend themselves effectively, making them more susceptible to fatal attacks compared to adults.

### 6.3. Temporal, Seasonal and Spatial Dynamics

#### 6.3.1. Seasonal Patterns and Behavioral Insights

The Uttarakhand Forest Department links most leopard attacks to monsoon months [Hindustan Times, 2015] due to dense undergrowth providing cover for leopards. However, this study highlights a distinct seasonal pattern: incidents peak in winter (n=121), followed by monsoon (n=88) and summer (n=81) (Table 11).

**Table 11.**
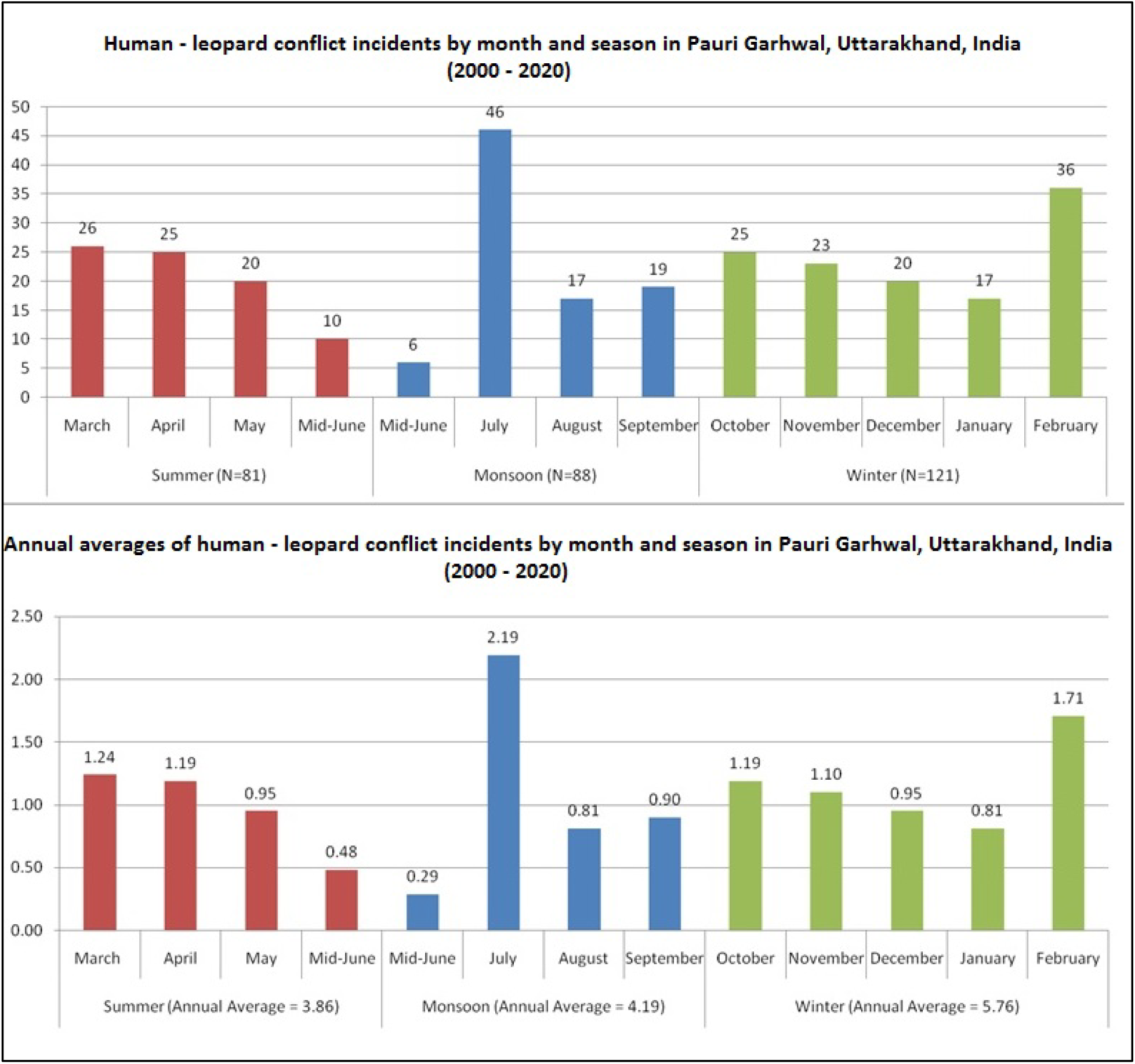
Human – leopard conflict by month and season in the Pauri Garhwal District, Uttarakhand, India (2000 – 2020)

Winter conditions—reduced daylight, extended nocturnal hours, and fog (Fig. 29a & 29b) — create an extended temporal window for conflict.

**Figure 29a & 29b.**
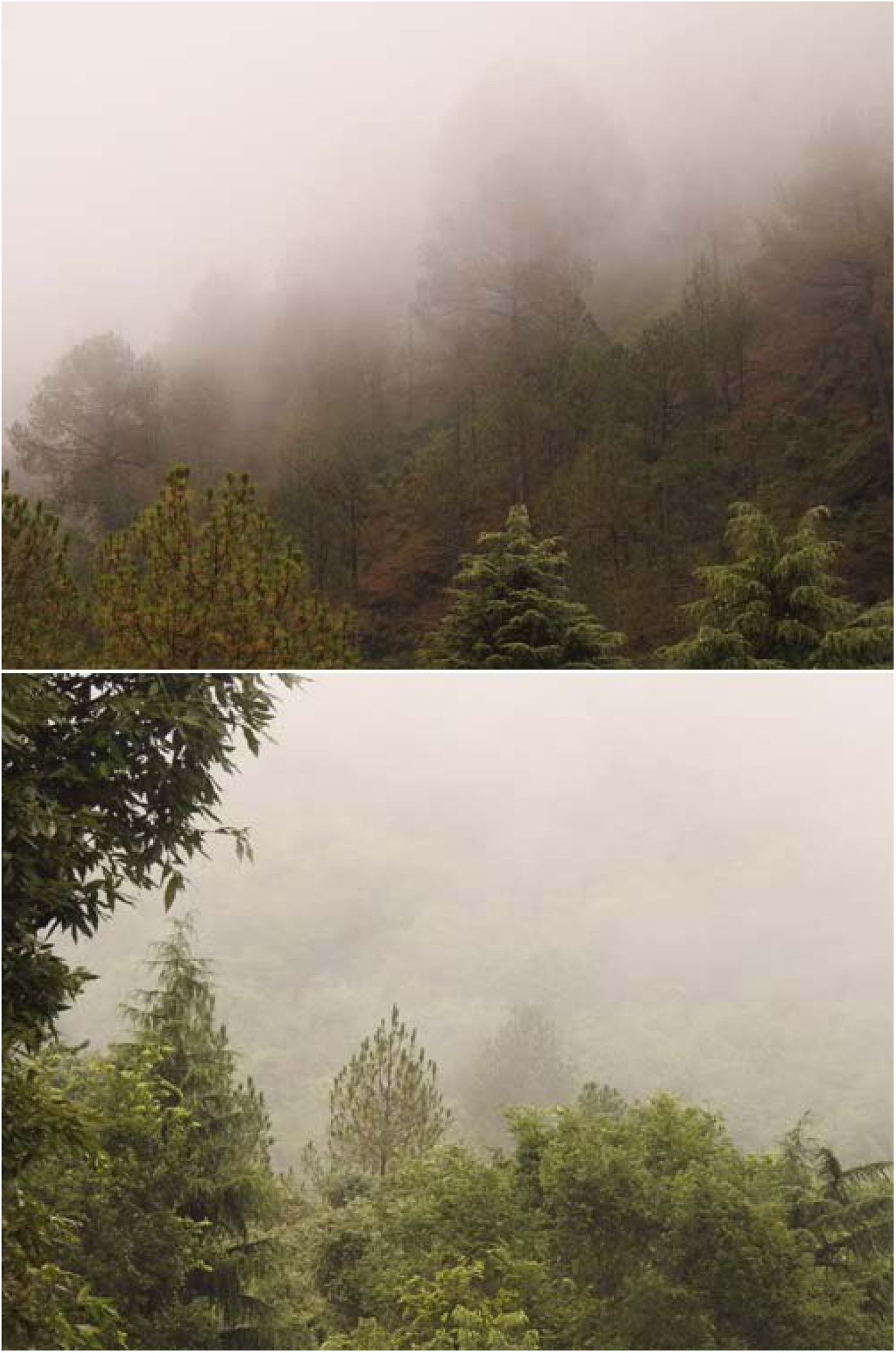
Winter fog in the Gadoli and Manda Khal Fee Simple Estates, Pauri Garhwal, Uttarakhand, India

Winter’s longer duration (151 days, 41.37% of the year) compared to summer and monsoon (107 days each, 29.32%) creates a greater temporal window for conflict, resulting in more incidents during winter than in summer or monsoon. Increased human activity in forests post-monsoon, including grazing, fuelwood collection, and commuting, elevates the risk. The monsoon months, with restricted movement in the mountainous terrain, create a scarcity of resources, prompting heightened forest use in winter to compensate.

In contrast, the spike in attacks during July, a monsoon month, is less likely tied to human forest entry and may instead align with leopard reproduction patterns in the region. Local knowledge suggests leopards in the region give birth in April/May (*pers. Obs.*), making their cubs around three months old by July. During this time, leopard mothers likely require additional prey to feed their cubs. As prey-depleted forests push leopards toward human settlements for food, this need may significantly contribute to the spike in leopard attacks on humans in July.

#### 6.3.2. Spatial Patterns with Seasonal and Temporal Insights

Leopard attacks occur year-round in human habitation, with no seasonal preference. Incidents in forests occur during daytime human activities, whereas night time incidents reflect leopard activity around settlements. Attack timings in human habitation (6:00–22:00) peak thrice daily, with most incidents between 16:00–21:00 suggesting leopards are most active during this period but remain near settlements, especially those close to forests, throughout the day. Daylight hours shorten from 06:00–18:30 in summer to 06:30–16:30 in winter. Shorter winter daylight, shifts attack timings earlier in human habitation. In forests, people typically enter at mid-day during summer and monsoon for activities like fuelwood collection and grazing but enter later in winter, shifting conflict incidents from mid-day in summer and monsoon to late afternoon and early evening in winter. Human nighttime presence in forests raises concerns of poaching or illegal activities including tree felling.

### 6.4. Effectiveness of Ecological Restoration

#### 6.4.1. Ecological Restoration and Lack of Conflict

Human-leopard conflict incidents in the Pauri sub-division and the district remained consistent from 2000–2020, ruling out a general decline in conflict incidents as the reason for zero incidents in the Gadoli and Manda Khal Fee Simple Estates. This absence of conflict is likely attributable to targeted conservation efforts promoting ecological restoration and ecosystem recovery, which have effectively prevented human-leopard interactions in these estates, through habitat preservation, legal action against illegal infrastructure development, and conservation-oriented land management. The comprehensive approach to habitat enhancement in the Gadoli and Manda Khal Fee Simple Estates has created a suitable environment for leopard prey, likely reducing stress on leopards compared to the broader landscape. Prey presence is monitored through visual sightings, sign surveys, and camera traps, which also track prey reproduction (Video File 7: https://doi.org/10.6084/m9.figshare.23620626.v1). Notably, prey species that were once secretive and nocturnal are now more visible during daylight hours (*pers. Obs.).* This improved prey presence and reduced leopard stress likely contribute to the absence of human-leopard conflict over two decades, particularly for an area that lies within an epicenter of such conflicts in the Pauri sub-division.

#### 6.4.2. Leopard Behaviour and Prey

Although no studies on leopard home ranges exist for Uttarakhand’s central Himalayas, research from Himachal Pradesh [Odden et al., 2014] indicates a minimum home range of 15 km². This suggests that leopards in the Gadoli and Manda Khal Fee Simple Estates share territories that overlap adjacent villages and forests. Despite this overlap and frequent attacks in peripheral areas, no incidents were recorded within these estates over two decades. It is hypothesized that habitat quality in the estates reduces leopard stress and encourages the presence of natural prey availability, decreasing the likelihood of conflict. However, further research into this is required.

#### 6.4.2. The Covid – 19 Lockdown and Conflict Resolution

During the 3-month global lockdown in 2020 (April-July), Pauri Garhwal enforced curfews implemented by the local police which restricted access to forests, effectively making them in-violate. No human – leopard conflict incidents occurred in 2020 during the lockdown. However, after restrictions were lifted in mid-July 2020, human-leopard conflict resumed with incidents corresponding to the re-entry of people into forests. This demonstrates that regulating human entry into forests can effectively reduce conflict as in the Gadoli and Manda Khal Fee Simple Estates and suggests that such measures, if applied on a larger scale, could help mitigate human-leopard conflict, particularly in forested regions.

### 6.5. Policy and Management Implications

#### 6.5.1. Alienation of *Van Panchayats*

Currently, the Uttarakhand Forest Department lacks habitat protection or improvement measures for community-managed forests, which face higher degradation rates than reserve forests [Personal Communication; Letter Number 1360/22-1 dated 21/11/2019, Garhwal Forest Division, Pauri, Uttarakhand Forest Department]. Legislative amendments from 1976 to 2012 diluted Van Panchayat laws, alienating communities from managing their forests [Naaz & Sahu, 2018; Singh, 2013]. This is further reinforced by amendments in 2024 (*pers. Obs.*) The 1931 legislation granted *Van Panchayats* autonomy to create and implement forest management plans without government interference. However, revisions have placed these customary rights under the preview of micro-plans devised by the Uttarakhand Forest Department, significantly reducing local decision-making power. This alienation has fostered ecologically unsound practices, diminished preservation efforts, and increased exploitation, contributing to deforestation and ecosystem decline. As outlined in Somanathan [1991], the absolute monopoly of the state has destroyed the incentive to use forests sustainably by the local communities in the Central Himalayas; where as comparatively in the mid – Himalayas of Nepal, forest cover which decreased by 0.96% annually between 1976 - 1991 increased by 0.63% annually from 1991 onwards after the intervention of the Community Forestry Management System [Tripathi et al., 2020].

#### 6.5.2. Impact of Infrastructure and Environmental Degradation

Linear infrastructure, like power lines and roads fragment forests and deplete wild herbivore populations. Power lines can trigger forest fires, while roads increase human access, intensifying pressures like grazing and fuelwood collection. Forest fires, often human-induced, unrelated to natural cycles, alter forest structure, soil properties, and moisture. Alongwith with climate change, recovering burn areas may result in a vegetation structure different from what occurred prior to the burn [Robinne, 2021]. Repeated burns often prevent forest recovery, leading to vegetation changes that drive prey species away and towards more favourable habitats [Smith, 2000]. Burn areas also increase predator visibility, reducing predation success [Smith, 2000].

Prey-depleted forests push leopards toward villages, where open garbage dumps (Fig. 30a & 30b) attract feral pigs, dogs, and cattle, increasing human-leopard encounters.

**Figure 30a & 30b.**
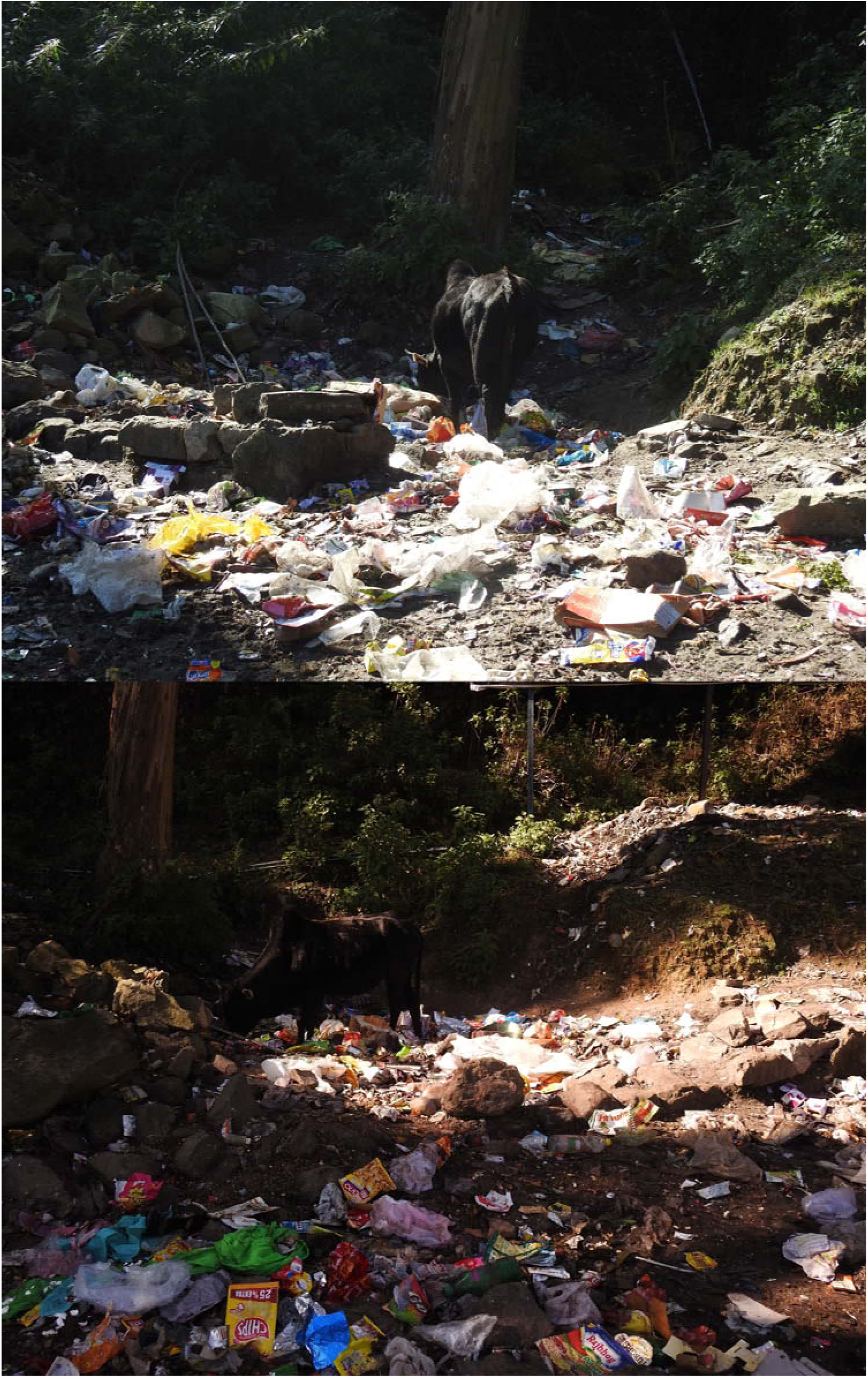
Open garbage dumps on a village-forest interface, Pauri Garhwal, Uttarakhand, India.

#### 6.5.3. Effectiveness of Conservation Efforts and Mitigation Projects

Several human-leopard conflict mitigation projects in Uttarakhand, such as the Indo-German Cooperation Project on HWC [Indo-German Human-Wildlife Conflict (HWC), 2021], National Mission on Himalayan Studies (NMHS) [Rawat & Sathyakumar, 2018], WCS-India in partnership with the Titli Trust and the Uttarakhand Forest Department [Sondhi et al., 2016], and the Wildlife Institute of India [Agarwal et al., 2011], focus on community engagement, conflict data analysis, GIS mapping, and awareness programs. Despite these efforts, human-leopard conflict remains unabated in the Pauri Garhwal district, as shown by this study. In contrast, conservation efforts aimed at ecological restoration in the Gadoli and Manda Khal Fee Simple Estates have resulted in zero human-leopard conflict.

#### 6.5.4. Ineffectiveness of Lethal Control

The Uttarakhand Forest Department primarily relies on lethal control for human-leopard conflict mitigation. Between 2006 and 2020, the removal of 324 leopards, averaging 21.6 per year, was sanctioned [Personal Communication; Letter Number 1051 / 22-1(7) / (27) dated 30/9/2020, Office of the Principal Chief Conservator of Forests (Wildlife), Dehradun, Uttarakhand Forest Department]. However, studies indicate that lethal control may exacerbate conflicts. For example, removing carnivores creates territorial vacancies, attracting transient individuals and increasing conflict rates [Herfindal et al., 2005; Harper et al., 2008; Athreya et al. 2011; Elbroch & Treves,2023]. Cases from South Africa and Maharashtra demonstrate similar outcomes, where leopard culling and translocation heightened conflicts [Conradie & Piesse, 2013; Athreya et al., 2011].

#### 6.5.5. Lack of leopard population monitoring and its implications

The Uttarakhand Forest Department lacks a program to monitor the leopard population in Pauri Garhwal, making it difficult to assess human-leopard conflict in relation to leopard population trends. Between 2000 and 2020, human-leopard conflict remained consistent. However, if the leopard population had increased during this period, it would suggest a reduction in conflict relative to leopard population growth. Conversely, a decline in the leopard population would indicate an increase in conflict despite fewer leopards.

#### 6.5.6. Underlying Conflict Dynamics

The IUCN SSC Guidelines on Human-Wildlife Conflict and Coexistence [IUCN, 2023] highlight that many conflict mitigation strategies fail to address underlying dynamics, often leading to short-lived or worsening solutions. Human-wildlife conflict typically arises from spatial and temporal overlap between wildlife and people, particularly when they share land, resources, or habitats. This interface is dynamic, and higher encounter rates increase the likelihood of conflicts. Factors driving this interface include changes in land use, human encroachment into wildlife habitats, and shifts in wildlife distribution, often linked to species’ dietary needs, which may bring them closer to human settlements. This is aptly portrayed in human – leopard conflict presently being witnessed in the district of Pauri Garhwal as outlined below.

In Pauri Garhwal, many human-leopard conflict incidents occur in forests which are exploited for fuelwood, fodder, illegal grazing, and illicit lopping, as well as being used for travel between villages by foot. These activities increase the likelihood of encounters between leopards and humans in forest areas. Infrastructure development, forest conversion, and encroachments further degrade habitats, causing deforestation and fragmentation. While leopards can adapt to habitat changes, their prey species cannot, leading to prey-depleted forests. Female leopards, struggling to secure food for themselves and their cubs, may venture closer to human settlements, resulting in increased conflicts. Garbage dumps attract cattle, feral pigs, and dogs, which in turn attract leopards to villages. Human encroachment and expanding settlements near forests increase opportunities for leopards to enter these human habitations. Unaccompanied children are particularly vulnerable, and once a leopard learns humans are easy prey, repeat attacks may occur.

Human-leopard conflict in Pauri Garhwal is driven by both top-down processes, such as government policies promoting infrastructure development, encroachments, and settlement expansion, and bottom-up processes, like man-made forest fires, illegal felling, and grazing [Hunter & Price, 1992; Gandiwa, 2013] in forest areas. These activities, influenced by population density, lead to deforestation and prey depletion, pushing leopards into closer contact with humans, most likely as a result of ecological cascades caused by these human actions.

## 7. Conclusion

Ecological restoration for human-leopard conflict mitigation has shown promising results in the Gadoli and Manda Khal Fee Simple Estates, marking a novel approach in the Pauri Garhwal district. By safeguarding forests, regulating their use, and promoting habitat recovery, ecological restoration offers a sustainable long-term solution, unlike lethal control or translocation methods. It improves habitat quality for leopard prey and reduces stress in leopards, leading to fewer conflicts. Reserve forests managed by the Forest Department can enhance prey availability, while community-managed *Van Panchayat* forests can provide resources for local communities and secure leopard habitats through sustainable management. Addressing government control over *Van Panchayats*, implementing the Forest Rights Act (2006) or involving the Indian Army’s “Eco –Task Force” of the Garhwal Regiment to work together with *Van Panchayats*, Civil Soyyam and private forests for habitat restoration and protection could strengthen community involvement and effective forest management.

Awareness programs for local communities on co-existing with leopards are essential for conflict mitigation. These should be paired with ecological studies and monitoring of leopard populations, prey, and habitats, supported by state agencies, conservation organizations, and private entities. Additional mitigation measures include establishing fuelwood and fodder plots with local native species, removing undergrowth near human settlements, improving garbage management, enforcing no-go policies for infrastructure in forests, removing encroachments, and implementing night lighting. Using camera traps, surveillance cameras, and bioacoustics to monitor leopard presence around human habitation should also be considered.

## 8. Study Limitations

While this study provides critical insights, several limitations warrant consideration. First, the reliance on secondary data sources, such as forest department records, may introduce biases or inaccuracies. Second, small sample sizes in specific analyses, limit the generalizability of findings. Additionally, while the study hypothesizes a link between *Van Panchayat* alienation and conflict escalation, direct evidence is lacking, necessitating further research to substantiate this claim.

## 9. Recommendations and Mitigation Strategies

To mitigate human-leopard conflict in Pauri Garhwal, an integrated approach, combining habitat conservation, community involvement, and policy measures is essential. Key strategies include:

- **Habitat Restoration**: Prioritize restoration in high-conflict areas, focusing on reforestation and creating wildlife corridors to connect fragmented habitats. Control forest fires and promote sustainable forest use.
- **Prey Base Recovery**: Enhance habitat restoration and quality to restore wild herbivore populations, supporting leopard prey availability, specially in high – conflict areas.
- **Research and Monitoring**: Conduct studies on leopard ecology and conflict patterns, using camera traps, bioacoustics, and GPS collars for real-time monitoring of movements in high-conflict zones.
- **Conflict Mitigation**: Establish protocols to identify and manage problematic leopards, minimizing reliance on lethal control.
- **Community Engagement**: Raise awareness about leopard behavior and promote non-lethal deterrents, such as night-time lighting.
- **Policy Development**: Regulate infrastructure to prevent habitat fragmentation and create conflict management policies balancing conservation with community welfare.
- **Knowledge Exchange**: Integrate successful practices from Africa, Latin America, and other parts of Asia.

By addressing these priorities, policymakers and conservationists can foster coexistence between humans and leopards while advancing broader conservation goals.

**Supporting Information:** https://doi.org/10.6084/m9.figshare.23620626.v1

Supplementary Material A: A-i and A - ii

1. Supplementary Material B: Detailed analysis and results, datasets and sub - datasets
2. Supplementary Material C: Time - Series

**Video Files:** https://doi.org/10.6084/m9.figshare.23620626.v1

## Manuscript Preparation

This manuscript benefited from OpenAI’s ChatGPT for language refinement, grammar checks, and structural improvements. The author independently reviewed and finalized the text, ensuring no generative AI was used in scientific data analyses and insights, interpretations, and conclusions which remained his own. The author reviewed and edited the content and takes full responsibility for the content of the publication.

**Ethical Approval:** This work did not require any specific permits or permissions.

**Funding Declaration:** This research did not receive any specific grant or any financial support from funding agencies or individuals in the private, public, commercial, or not-for-profit sectors.

## Acknowledgements

I, thank *The Gadoli and Manda Khal Wildlife Conservation Trust* for supporting this study as well as my various on – going research and conservation activities on leopards, leopard prey and their habitats in the private forests of the Gadoli and Manda Khal Fee Simple Estates, District Pauri Garhwal, Uttarakhand, India through the Leopard Ecology Project. I would also like to thank *Dr. Adriana Consorte-McCrea, PhD.* and *Jacob Owens, PhD.* for providing valuable suggestions that helped in improving this publication. This research did not receive any specific grant from funding agencies in the public, commercial, or not-for-profit sectors.

## Notes

https://doi.org/10.6084/m9.figshare.23620626.v1

### Competing Interest Statement

The authors have declared no competing interest.

### Summary of Updates

The revision has been submitted to correct the text, figures, and tables

